# DNA adduct and mutational profiles reveal a threshold of cellular defenses against *N*-nitrosodimethylamine administered to mice in drinking water

**DOI:** 10.1101/2025.11.13.687536

**Authors:** Nina E. Gubina, Lindsay B. Volk, Anna F. Dormitzer, Emily M. Michelsen, Lee J. Pribyl, Joshua J. Corrigan, Esha D. Dalvie, Amanda L. Armijo, Monét Norales, Nicolette A. Bugher, Kayla M. Schonvisky, Desiree L. Plata, Bevin P. Engelward, Robert G. Croy, John M. Essigmann, Bogdan I. Fedeles

## Abstract

*N*-Nitrosodimethylamine (NDMA) is classified as an animal and probable human carcinogen. Murine liver DNA adducts, mutations, MGMT and CYP2E1 were evaluated following chronic administration of NDMA in drinking water. In a dose-escalation study, 7-methylguanine (m7G) increased linearly with NDMA dose. *O*^6^-Methylguanine (m6G) remained near background for NDMA doses up to ∼1 ppm, beyond which its level, and corresponding mutations, rose steeply. An extended study was done with 5 ppm NDMA, in which adducts were measured at 3 weeks and mutations at 10 weeks. While the level of CYP2E1 was unchanged, MGMT gene transcription was induced in females at 10 weeks. Homologous recombination-mediated chromosomal rearrangements did not increase over background. Point mutations, however, were elevated substantially in both sexes. Mutational analysis over 96 trinucleotide contexts revealed predominantly GC→AT mutations in 5’-purine-G-3’ contexts in a pattern matching human COSMIC cancer mutational signature SBS11, with secondary features resembling SBS119 (AT→GC). Taken together, the data implicate m6G as the dominant mutagenic adduct under chronic dosing with NDMA. Furthermore, the genomic m6G level was identified at which its dedicated repair protein, MGMT, became saturated. The coordinated application of DNA adduct, mutational and biochemical analyses provides a new approach for early detection and cancer management.

**Graphical abstract:** 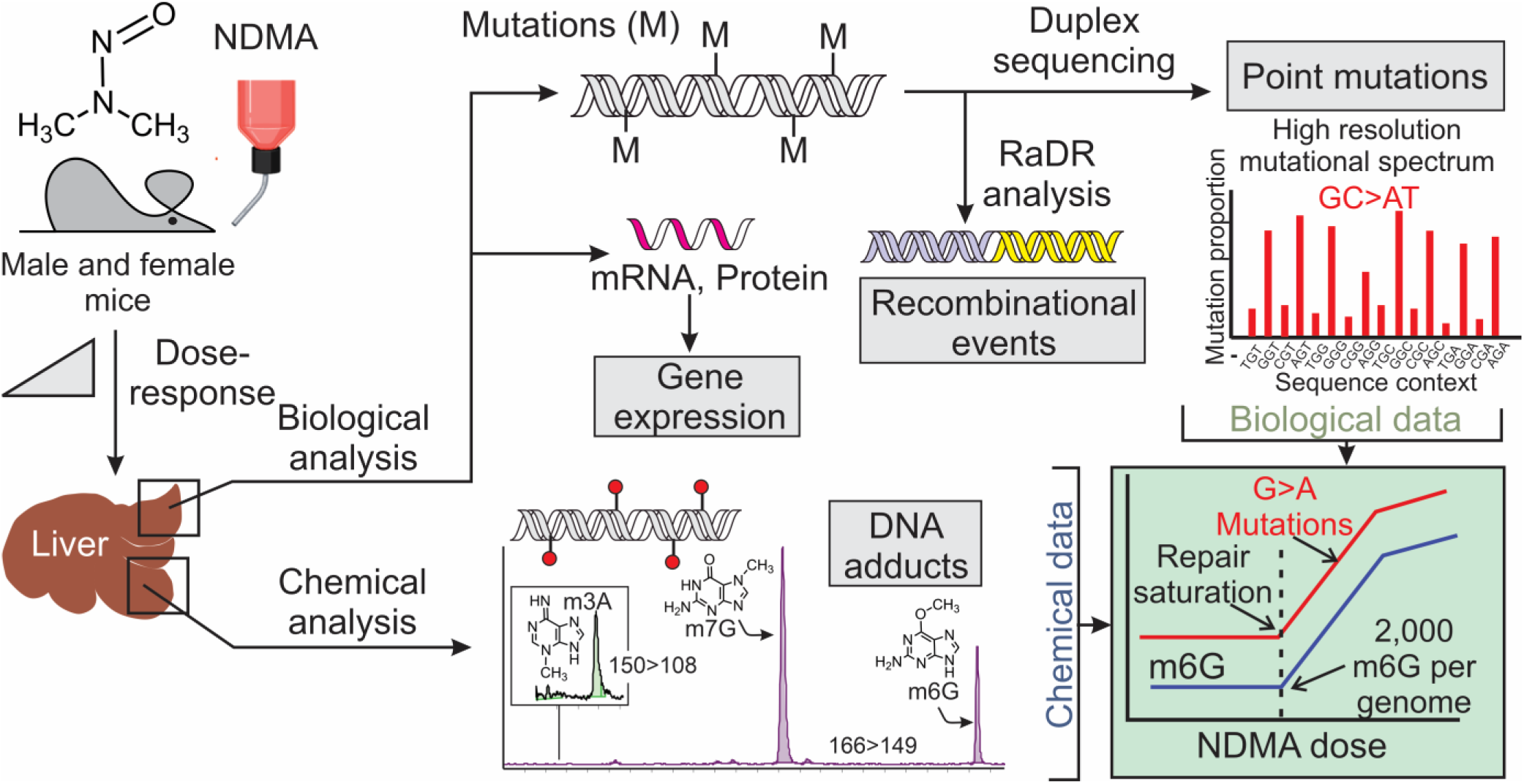

## Introduction

*N*-Nitrosodimethylamine (NDMA; Figure 1) is a member of the class of toxicants known as *N*-nitroso compounds (1), which are extraordinarily versatile carcinogens (2). In animals, NDMA primarily causes liver, respiratory tract, blood vessel, bile duct and kidney cancers, and, in humans, exposures are linked to the risk of liver, stomach, esophageal, colorectal, bladder and prostate malignancies (2–4). The International Agency for Research on Cancer lists NDMA in Group 2A as a probable human carcinogen (2). People are exposed to NDMA from a wide variety of sources (5). It is present in food, especially nitrite-cured meats and fish (5–7); it appears as a byproduct of industrial chemical processes (5, 8, 9); it is found in and near some Superfund sites (10); it forms in water treated by chlorination or chloramination to prevent infectious diseases (11, 12); it is present along with other *N*-nitroso compounds in tobacco products (13); and it is a contaminant arising from the production of certain drugs, such as valsartan, metformin and ranitidine (14–16). In addition, NDMA is formed naturally at low levels in animals including humans, presumably from activation of the innate immune system (17–20). While humans certainly are exposed to NDMA, it is noteworthy that the amounts of exposure from the aforementioned scenarios vary by orders of magnitude. Hence, uncertainty exists regarding whether the routes or the extents of exposure are sufficient to trigger disease. This question is especially important because most organisms have inherent or inducible protective factors that can mitigate disease risk (21–24).

**Figure 1.**
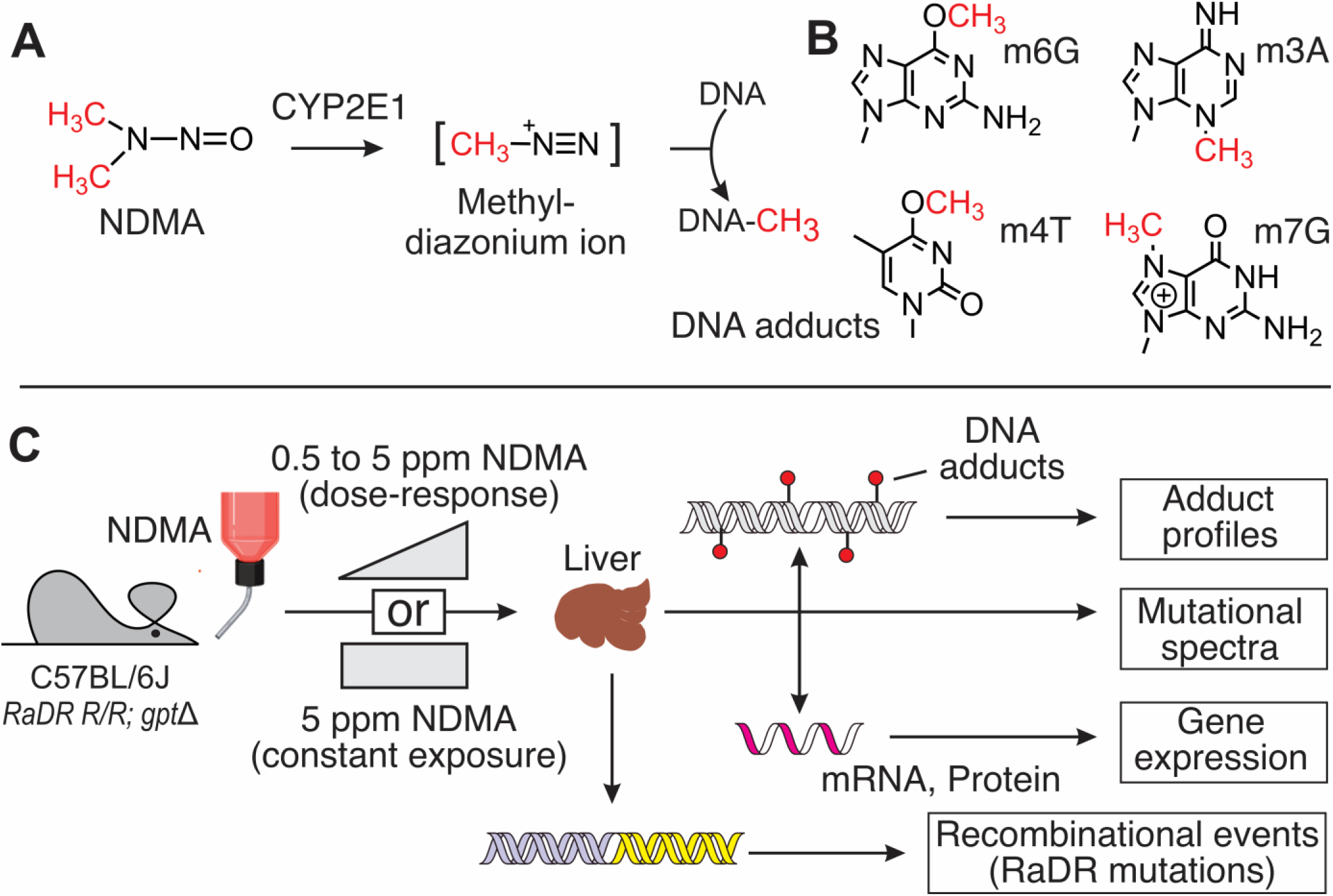
Experimental design and endpoints. **A,B.** *N*-Nitrosodimethylamine (NDMA) is metabolized by cytochrome P450 2E1 yielding methyldiazonium, a reactive intermediate that methylates DNA to form a wide range of DNA adducts. The most abundant or biologically consequential adducts are: 7-methylguanine (m7G), 3-methyladenine (m3A), *O*^6^-methylguanine (m6G) and *O*^4^-methylthymine (m4T). **C.** In the present study, NDMA was administered to mice in drinking water at concentrations ranging from 0.5 ppm to 5 ppm, as indicated. Liver DNA, RNA and protein were isolated and analyzed for DNA adducts, mutations and protein expression of cytochrome P450 2E1 and the DNA repair protein MGMT.

While there are no evidence-based US federal drinking water standards for NDMA, the World Health Organization has issued a guideline for drinking water at 0.1 μg/L (0.1 ppb) (25). A number of laboratory studies over the past 70 years have examined the toxicity and carcinogenicity of NDMA at various doses in animal, primarily rodent, models (3, 18, 26). Because human exposure can occur from contaminated water, some of these studies have examined the dose-response relationships between chronic exposure to NDMA in drinking water and carcinogenesis in rodents, predominantly rats (27, 28). A smaller number of studies have been done in mice (29–32), with a few coming out only very recently (33, 34), despite the fact that this species is highly amenable to genetic analysis and manipulation (31).

The biochemical and genetic mechanisms by which NDMA causes the conversion of normal cells to cancer cells have been the subject of intense historical scrutiny. In the liver or other target organs, cytochrome P450 2E1 (CYP2E1) (35) aids in the conversion of the parent compound into a putative methyldiazonium ion, which then alkylates nucleophilic centers on the nucleobases and phosphates of DNA (Figure 1A). 7-Methylguanine (m7G), 3-methyladenine (m3A) and *O*^6^-methylguanine (m6G) are the quantitatively most abundant adducts, and the latter two (m3A and m6G) are widely believed to be the most biologically significant (Figure 1B). Site-specific placement of m6G in the genome of a virus, which was then replicated *in vivo*, revealed that this DNA adduct pairs with thymine during replication causing G→A mutations (36, 37), which are explained by structural (38, 39) and biochemical (40, 41) studies *in vitro*. The mutagenicity of m6G *in vivo* is dependent upon its local sequence context (42), incentivizing studies, such as the one presented here, in which a high-resolution sequencing tool is used to reveal the way flanking nucleotides affect the mutagenic properties of DNA adducts.

Many studies on carcinogenesis are conducted under less-than-ideal circumstances, such as the use of large, acute doses of toxicant that might skew results by quickly saturating cellular defenses, such as DNA repair systems. Because humans are often exposed chronically and at low levels, the present study was designed to examine the formation of DNA adducts and the genetic effects of NDMA when given orally in drinking water (Figure 1C). A dose-escalation study was done to find the point at which the mutation frequency at a given dose inflects sharply upward reflecting, presumably, the point at which the capacity of the principal DNA repair protein, *O*^6^-methylguanine DNA methyltransferase (MGMT), which protects the genome from mutagenic and toxic alkylation damage, became saturated. Genetic studies at incremental doses of NDMA revealed a distinctive mutational pattern through high-resolution duplex DNA sequencing. In parallel with these genetic studies, DNA adducts were measured using a stable isotope dilution mass spectrometry method that was sufficiently sensitive to detect m6G, m7G and m3A at background levels, and higher, in the mouse genome. The results show that m6G levels track closely with mutation frequency from exposure to NDMA.

## Materials and methods

### Animals

Wild-type mice on a C57BL/6J genetic background were used. The engineered mouse strain used carried two genetic cassettes: a *Rosa26* direct repeat GFP cassette *(RaDR^R/R^)* and a *gpt* delta cassette *(gpt^g/g^)* (31, 43). Mice were housed in an AAALAC-accredited facility with procedures approved by the MIT Committee on Animal Care and in compliance with NIH guidelines. Upon arrival, they were provided normal chow (irradiated pelleted diet Isopro RMH 3000; Purina Mills, Inc., St. Louis, MO) and filtered water *ad libitum*. Upon weaning, male and female offspring were randomly designated for control or NDMA treatment groups (Table S1). Thirty-day-old mice were transferred within one week to purified AIN-93G diet (TD.00447; Envigo Teklad Diets, Madison, WI) and kept on this diet for the duration of the experiments. This formulation allows for ingredient control, does not require irradiation and can be prepared without antioxidants such as tertiary butylhydroquinone. AIN-93G diet was shaped as 0.5-inch pellets and air-dried. To preserve freshness in the absence of antioxidants, food pellets were divided into one kg batches, vacuum packaged and stored away from light at −20°C. Weekly portions were defrosted and kept at 4°C prior to use. AIN-93G food was replaced in the feeders twice a week. Mice were weighed Monday-Friday during the first two weeks of each experiment and, after that, every week until the end of the experiment (Table S2). Mice were euthanized after two, three and ten weeks of exposure to NDMA by CO_2_ inhalation (Table S1). Tissues were flash-frozen immediately after extraction and weighed before use (Table S3).

Mice were free of the following murine pathogens: ecto- and endoparasites, mouse parvovirus, mouse hepatitis virus, mouse rotavirus, minute virus of mice, Ectromelia, Sendai virus, pneumonia virus of mice, reovirus, theilovirus, lymphocytic choriomeningitis virus, *Mycoplasma pulmonis*, *Filobacterium rodentium*, polyoma virus and mouse adenovirus. Pathogen status was verified by full serological testing using sentinel mice every six months, and with an abbreviated serology panel every two months.

### NDMA

NDMA was synthesized in-house and characterized by the method reported by He *et al*. (44). Sealed ampoules were stored at −20°C. Prior to all experiments, a working solution of 2 mg/mL NDMA was prepared in Milli-Q water, aliquoted, and stored in light-protected tubes at −20°C with a single freeze-thaw cycle. NDMA is sensitive to light, but given the absorption peaks are in the UV range (227 and 332 nm), the risk of significant photodegradation under typical indoor lighting is generally low. Nevertheless, NDMA stability was tested under the conditions used for the dosing of mice. Using a formulation of 5 ppm, the stability of NDMA was evaluated under vivarium conditions (12 h light / 12 h dark cycling). Standard micro-isolator cages with autoclaved hardwood chip bedding were assembled with either red hazmat bottles (intended for use with NDMA) or with clear bottles covered in foil (control conditions in which 100% of light is blocked to ensure NDMA integrity). Water samples were extracted daily and analyzed for NDMA using an Agilent 7890B gas chromatograph with a 5977B mass spectrometer and chemical ionization (Agilent Technologies, Palo Alto, CA) following EPA Method 521 (45). It was found that 5 ppm NDMA in a standard red plastic water bottle is stable for three days (Figure S1; Table S4); consequently, bottles with NDMA solution were freshly prepared and replaced every two to three days. All operations with NDMA under the conditions of vivarium were performed following MIT Environment, Health & Safety Office and Committee on Animal Care guidelines.

### Dosing regimens

Three dosing regimens were used. Regimen A (Figure 2A) was a range-finding study in which DNA adducts and high-resolution mutational spectra were evaluated at the end of a two-week interval of dosing at 0, 0.5, 1, 2.5 and 5 ppm NDMA. For this experiment, four males were dosed at 0 and 0.5 ppm, and three males were dosed at 1, 2.5 and 5 ppm. Dosing occurred during the 6th and 7th week of life. Regimen B examined body weight (Table S2), DNA adducts (Table S5), CYP2E1 and MGMT mRNA and protein levels (Tables S6 and S7) after three weeks of dosing at 5 ppm NDMA, again starting at six weeks of age. Control mice received untreated water. Four males and four females per group were used in Regimen B (Figure 2B). Regimen C continued the conditions of Regimen B (5 ppm NDMA) for another seven weeks, ending at the 16th week of life (Figure 2C). Six males and six females per group were evaluated for body weight (Figure S2 A and B, Table S2), liver, lungs and kidneys weights (Figure S2 C, D, E; Table S3), for CYP2E1 and MGMT mRNA (Table S6) and protein levels (Table S7) in livers. Fresh left liver lobes were assessed for *RaDR* foci, which represent large scale gene rearrangements (Table S8). Livers from all experiments were flash-frozen in liquid nitrogen and stored at −80°C for further analysis. DNA isolated from the livers of mice on Regimen C were analyzed to determine high-resolution mutational spectra (HRMS) (Table S9-S10) and mutation frequency in the exogenous *gpt* gene (Table S11).

**Figure 2.**
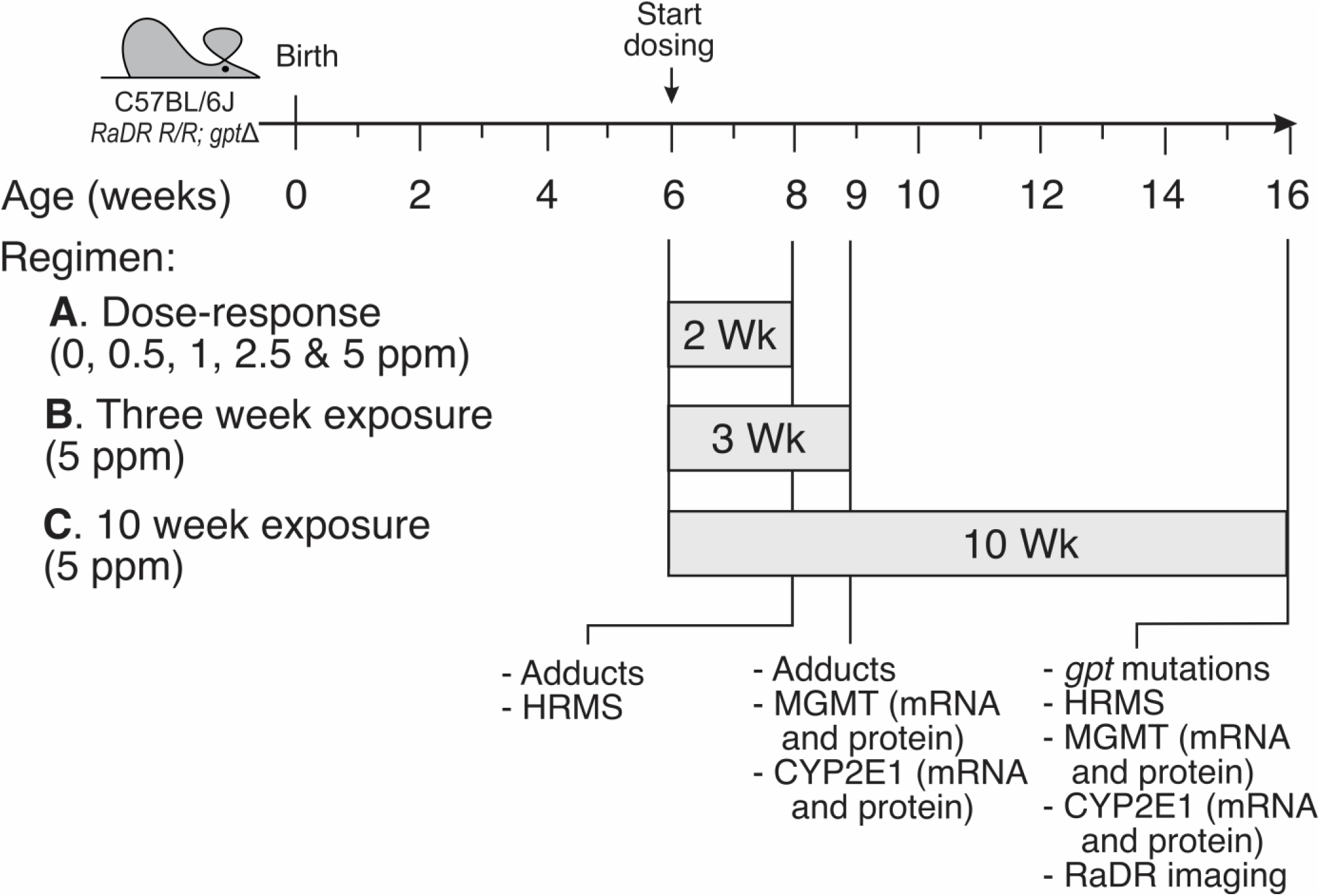
The dosing regimens for the animal studies described. Dosing of mice with NDMA or control water started at 6 weeks after birth and continued following three schedules. **A.** Dose escalation study involving exposure to four NDMA doses and control water for two weeks. Endpoints include measuring DNA adducts and high-resolution mutational spectra (HRMS) in liver. **B.** Exposure to 5 ppm NDMA for three weeks. Endpoints include measuring DNA adducts and mRNA and protein levels of CYP2E1 and MGMT. **C**. Exposure to 5 ppm NDMA for 10 weeks. Endpoints: mutation frequency by *gpt* assay, HRMS, mRNA and protein levels of CYP2E1 and MGMT, and *RaDR* imaging.

### Methods for determining DNA adducts in tissues

#### Preparation of methylated ^15^N-labeled *Escherichia coli* DNA (Figure 3A)

Isotopically labeled DNA was obtained from *E. coli* 15224 (ATCC) grown in M9 minimal medium containing ^15^N-ammonium chloride from Cambridge Isotope Labs (46). Two mL of a lysozyme solution (10 mg/mL in 0.25 M Tris buffer pH 8.0) were added to a frozen pellet of *E. coli* (5 g). After thawing, the cell suspension was placed on ice for 45 min. Cells were lysed by addition of an equal volume of 20 mM Tris buffer pH 8.0, 0.2 M NaCl, 20 mM EDTA, 1% SDS, containing 20 mg/mL proteinase K and incubated overnight at 37°C. The solution was then extracted with an equal volume of phenol:chloroform:isoamylalcohol (24:25:1). The aqueous phase was isolated via centrifugation and extracted a second time with an equal volume of chloroform following centrifugation. DNA was precipitated from the aqueous solution by addition of 1/10 volume of 3 M ammonium acetate pH 5.3 and 2.5 volumes of cold ethanol and was collected on a glass rod. After washing with 70% ethanol, the DNA was briefly dried, dissolved in TE buffer (pH 7.5) and treated with RNase A (100 μg/mL) for 2 h at 37°C. The solution was then subjected to a second round of phenol/chloroform/isoamyl alcohol extraction, after which the DNA was precipitated with ethanol as described above.

**Figure 3.**
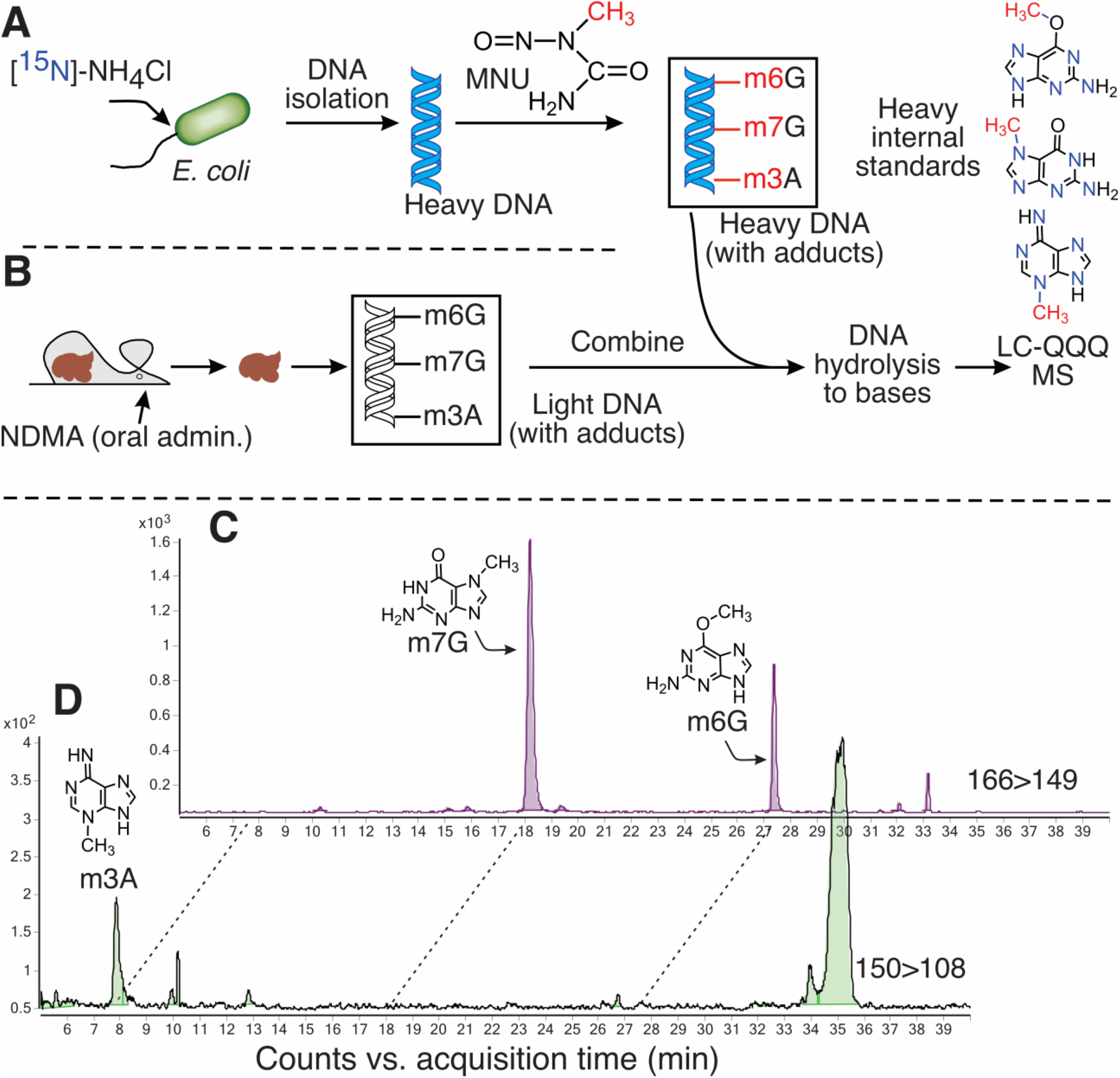
DNA adduct analysis overview. **A**. Heavy nitrogen (^15^N)-labeled ammonium chloride was fed to *E. coli* bacterial cells growing in minimal media to obtain pan-labeled heavy DNA. The heavy DNA was subsequently treated with MNU, to generate ^15^N-labeled methylated DNA adducts. The heavy DNA containing adducts was then used to provide internal standards for the LC-QQQ-MS analysis. **B**. Mice were exposed to NDMA via oral administration through drinking water. After 2-week or 3-week exposures, the mice were sacrificed, DNA from their livers was isolated, hydrolyzed, mixed with heavy DNA internal standards and subjected to the quantitative adduct analysis by LC QQQ-MS. **C**. Typical chromatography trace demonstrating separation of 7-methylguanine (m7G) and O6-methylguanine (m6G) adducts when monitoring a characteristic methylated guanine mass transition at 166>149. **D**. Typical chromatography trace indicating the elution time of 3-methyladenine (m3A) when monitoring its characteristic mass transition at 150>108.

Two mg of the ^15^N-labeled DNA was allowed to react with *N*-methyl-*N*-nitrosourea (MNU; 500 μM) in aqueous solution for 1 h, after which the DNA was precipitated with 3 volumes of cold ethanol in the presence of 0.3 M ammonium acetate pH 5.3. The methylated ^15^N-DNA was washed with 70% ethanol, dried, and dissolved in 2 mL of purified water. After having been dissolved in DNAse/RNAse free water at ∼1 mg/mL, aliquots of MNU-modified ^15^N-labeled DNA were stored at −20°C.

#### Isolation of DNA from frozen liver tissue, DNA hydrolysis and solid phase extraction (Figure 3B)

Adduct formation was evaluated in livers of mice treated with NDMA/control water for two (Regimen A) or three (Regimen B) weeks. DNA was isolated from 100 mg of frozen liver tissue using Qiagen 100/G Genomic tips (Qiagen, Germantown, MD, Cat# 10243) following the procedure recommended for genomic DNA preparation. Briefly, liver tissue was homogenized in buffer QBT in the presence of RNase and proteinase K and incubated overnight at room temperature. The solution was then loaded onto a Qiagen 100/G tip, washed with buffer QC, and retained DNA was eluted with buffer QF. DNA was precipitated with isopropanol and recovered by spooling onto a glass rod, then subsequently washed with 70% ethanol, briefly dried and then dissolved in 500 μL of purified water. The amount of recovered DNA was determined via UV absorption using a NanoDrop One C spectrometer (ThermoScientific, Waltham, MA, USA).

DNA (100-125 μg) containing 10 μg of methylated ^15^N-DNA, which served as an internal standard, was hydrolyzed by addition of 50 μL 90% formic acid and heating at 95°C for 1 h. Hydrolyzed DNA samples containing isotopically labeled standards were loaded onto Strata X SC cartridges (100 mg; Phenomenex, Torrance, CA), which had been prewashed with acetonitrile followed by 0.1 N formic acid. The cartridges were then washed with 6 mL of 0.1 N formic acid. The retained compounds were eluted with 1.5 mL of a solution containing 5% ammonium hydroxide, 10% ethanol and DNAse/RNAse free water, and vacuum centrifuged to dryness at room temperature. Samples were reconstituted in 100 μL of 0.1 N HCl and transferred to glass vials for LC/QQQ MS analysis.

#### LC/QQQ MS analysis of DNA adducts (Figure 3C&D)

LC/ESI-MS/MS analysis was performed on an Agilent 6410 Triple Quadrupole mass spectrometer interfaced with an Agilent 110 series HPLC (Agilent Technologies, Palo Alto, CA). Samples (25 μL injected) were resolved on a 4.6 x 100 mm, 2.7 μm particle size InfinityLab Poroshell column (Agilent Technologies, Palo Alto, CA) and eluted at a flow rate of 0.5 mL/min with a step gradient of 4 mM ammonium acetate containing 2% acetonitrile (solvent A) to 100% acetonitrile/0.0001% acetic acid (solvent B) over a 40 min period followed by column re-equilibration. QQQ MS analysis was performed in the positive ion mode using N_2_ as the nebulizing/drying gas. Capillary voltage and temperature were set 4.0 kV and 350°C, respectively. Selective reaction monitoring was used for quantitative analysis of methylated purines with the collision energy set at 25 for methylated guanines and 30 for the methylated adenine. Nitrogen was used as the collision gas. The following transitions were monitored: m/z 166 [MH^+^] → m/z 149 for m6G and m7G (Figure 3C); m/z 171 [MH^+^] → m/z 153 for *O*^6^-methyl-^15^N-guanine and 7-methyl-^15^N-guanine; m/z 150 [MH^+^] → m/z 108 for m3A (Figure 3D) and m/z 155 → m/z 112 for 3-methyl-^15^N-adenine. The results of adduct analysis are presented in Table S5.

### *Gpt* delta assay: detection of point mutations in the *gpt* reporter gene in the *gpt* delta mouse genome

The *gpt* delta assay is a technology using transgenic *gpt* delta mice to probe small mutations, such as single nucleotide substitutions, small insertions, deletions and frameshifts (43). *Gpt* delta mice carry a tandem array of more than 50 copies of the 48 kb λ-EG10 phage genome on chromosome 17. Each phage genome includes a 6.4 kb insert, which is later converted to a plasmid containing the 459 bp *E. coli guanine phosphoribosyltransferase* gene (*gpt*) and the chloramphenicol acetyltransferase selection marker. The phage are rescued from DNA via a lambda *in vitro* packaging reaction. For the *gpt* delta assay in this study, we used a protocol adapted from Armijo *et al.* (31). Specifically, livers were collected after 10 weeks of treatment with control/NDMA in drinking water (n=6 for each group), flash frozen in liquid nitrogen and stored at −80°C until further analysis. Genomic DNA was extracted from ∼40 mg of liver tissue using the RecoverEase DNA Isolation Kit (Agilent Technologies, Santa Clara, CA, cat# 720202). The λ-EG10 phage was packaged *in vitro* with genomic DNA using a Transpack packaging extract (Agilent Technologies, Santa Clara, CA, cat# 200220). The quality of packaging reactions was confirmed by growing the phage in *E. coli* XL-1 Blue MRA cells. Samples with good quality of packaged phage (one million PFU or more) were transfected into an *E. coli* YG6020 bacterial strain expressing the *cre* gene (43) and containing a 6.4 kb plasmid that included the *gpt* gene. The full genotype of XL-1 Blue MRA cells: (*Δ(mcrA)183 Δ(mcrCB-hsdSMR-mrr)173 endA1 supE44 thi-1 gyrA96 relA1 lac*). The full genotype of the YG6020 strain: (*F-, araDl39, A(ara, liu)7697, AlacX74, gaW, galK, rpsL, deoR, qBOdlacZhM15, endAI, nupC. recAl, mcrA, A(rnrr, hsdRMS, rncrBC, Δ(gpt-proA)62 and cre+* (43).These bacteria were cultured on selective media containing chloramphenicol (CM, 25 μg/mL, Sigma-Aldrich, St. Louis, MO, cat# 501957627) and 6-thioguanine (6-TG, 25 μg/mL, Sigma-Aldrich, St. Louis, MO; cat# A4882-1G) or CM alone. 6-TG resistance was confirmed by regrowth of colonies on plates containing CM and 6-TG. The mutant frequency (MF) per million colonies for each group was calculated as a ratio of the total number of 6-TG-resistant colonies to the average number of CM-resistant colonies, accounting for the dilution factor, and multiplied by one million: MF_per_million_=6TG_total_/(CM_average_x6000) x 10^6^. The results of the *gpt* assay are shown in Table S11.

### *RaDR* assay: determination of large chromosomal rearrangements in the *RaDR* locus

Livers were collected after 10 weeks of treatment with 5 ppm NDMA in drinking water, with toxicant-free water as a control (n=6 for each group). Freshly excised left liver lobes were held on ice in 0.01% trypsin inhibitor (Boston BioProducts, Milford, MA, cat# P-1540) dissolved in PBS prior to imaging. The entire left lobe of the liver was secured between a glass slide and a cover slip. The imaging and data analysis were performed as described by Kay *et al.* (47). Specifically, the dorsal surface of each liver was imaged with a Nikon Eclipse Ti2 scanning microscope (Nikon Metrology Inc., Brighton, MI) on the 2x objective in the FITC channel with emission wavelength 535 nm using an Andor Zyla 4.2 camera (Oxford instruments, Concord, MA) and NIS Elements software. The resultant raw .tiff images were submitted to analysis by a user-trained two-stage machine learning algorithm, which was used to identify and enumerate fluorescent foci within intact tissue (48). The representative *RaDR* images for the control and NDMA-treated mice are shown in Figure S3. All liver images were analyzed by the machine learning program based on parameters developed from training data (for the details, see Kay *et al.* (47)). The number of fluorescent foci was normalized to the area of the liver in the image (Table S8). After imaging, livers were embedded in Tissue-Tek O.C.T. compound (Sakura Finetek USA, Inc., Torrance, CA), flash-frozen in liquid nitrogen and stored at −80°C prior to further use.

### Duplex consensus sequencing of DNA

Duplex DNA sequencing, which is four orders of magnitude more accurate than conventional next-generation sequencing (NGS) (49), provides a high-resolution fingerprint of mutations across targeted areas of the genome. High molecular weight DNA was isolated from ∼10 mg of liver using the Monarch Genomic DNA Purification Kit (New England Biolabs, Ipswich, MA, Cat# T3010S) following instructions from the manufacturer. DNA was eluted in 50 μL of elution buffer, and DNA concentrations were measured by a NanoDrop OneC (ThermoScientific, Waltham, MA, USA) (Table S9). Samples were kept at 4°C prior to use. Approximately 1000 ng of genomic DNA was used to prepare a sequencing library using the TwinStrand DuplexSeq Kit (TwinStrand Biosciences, Inc., Seattle, WA) following instructions from the manufacturer. DNA concentrations at the quality control steps were measured by a Qubit 4 Fluorometer (Invitrogen, Carlsbad, CA, cat# Q33226) using Qubit TM 1X dsDNA BR Working Solution (Invitrogen, Carlsbad, CA, cat# Q33266) (Table S9). Samples were then submitted for NGS at the MIT BioMicro Center core facility (https://bmcwiki.mit.edu/index.php/BioMicroCenter). Quality control performed on the AATI Advanced Analytical Fragment Analyzer (Agilent Technologies, Palo Alto, CA) showed an average library fragment size of 400 bp. Libraries were prepared using a low volume pipetting robot mosquito HV genomics (SPT Labtech, Melbourn, United Kingdom), and Illumina TruSeq Library Prep (Illumina,San Diego, CA, cat# 20015963) was completed following the instructions of the manufacturer. Samples were pooled and submitted for sequencing on a NovaSeq 6000 instrument (Illumina,San Diego, CA, cat# 20012850) in one lane of an S4 flow cell with 150 + 150 paired-end, v2 chemistry.

### Next-generation sequencing data analysis

The raw data files generated from sequencing (fastq) were compressed (gzip) and then processed using the TwinStrand DuplexSeq Mutagenesis App (v4.5.0) running on the DNANexus platform. The App generates mutation-position files (.mut) that enumerate the point mutations for each sample. Lists of unique mutations, where each change at a given genomic location is counted only once, were constructed and then plotted as trinucleotide mutational spectra, as previously described (31). The trinucleotide mutational plots show mutation frequencies per-trinucleotide, which were obtained by normalizing mutational frequencies by the frequencies of appearance of each trinucleotide in the genomic target under analysis. Additionally, comparisons among experimental spectra and among COSMIC mutational signatures (v3.4 repository) were performed using cosine similarity metric, as previously described (31).

### Measurement of gene expression

Livers were collected after three and ten weeks of exposure to NDMA in drinking water (n=4 for three weeks, n=5 for 10 weeks), flash frozen in liquid nitrogen, and stored at −80°C until further analysis. RNA was isolated from ∼20 mg of liver using RNEasy Mini Kit (Qiagen, Germantown, MD, Cat # 74104) and treated by the RNase-Free DNase Set (Qiagen, Germantown, MD, Cat# 79254) following instructions from the manufacturer. RNA elution was performed twice with the same 40 μL of nuclease-free water, and its purity and concentration were measured using the NanoDrop OneC instrument (ThermoScientific, Waltham, MA, USA) (Table S6).

Synthesis of first-strand cDNA was performed on 1 μg RNA in 20 μL of incubation mixture using LunaScript® RT Master Mix Kit (Primer-free) (New England Biolabs, Ipswich, MA Cat# E3025S) with 6 μM Random Primer Mix (New England Biolabs, Ipswich, MA Cat# S1330S) following the instructions from the manufacturer. The program for cDNA synthesis included the following steps: 25°C for 2 min (pre-anneal); 55°C for 10 min (cDNA synthesis); 95°C for 1 min (heat inactivation). Then, the cDNA was cooled down to 4°C and stored at −20°C prior to use. QPCR was performed on a LightCycler 480 Real-Time PCR System (Roche Diagnostics, Rotkreuz, Switzerland, BioMicro Center at MIT) using SYBR Green I-based Luna® Universal qPCR Master Mix (New England Biolabs, Ipswich, MA Cat# M3003L, no ROX) following the manufacturer’s protocol. The amount of each primer was 250 nmol. For each gene, the difference among samples was determined based on the calibration curve composed of four 10-fold dilutions of the 2.5-fold diluted equimolar pool of all cDNAs. cDNA for each sample was diluted 10-fold for the analysis to ensure all samples stayed within the range of the calibration curve. For calibration dilution standards and samples, 2 μL were added to 18 μL of qPCR incubation mix. Each sample and standard were assayed in duplicate in a clear 96-well plate. Two wells for each pair of primers were used as blanks, with DNAse/RNAse free water as samples. The program for all genes was the same, except for the temperature of annealing, and included initial denaturation for 60 sec at 95°C and 40 cycles of the following steps: 95°C for 5 sec; temperature of annealing for 20 sec; 72°C for 30 sec. Fluorescence was read at the final step of each cycle. Primers for qPCR with respective temperatures of annealing and product lengths are summarized in Table 1.

**Table 1.**
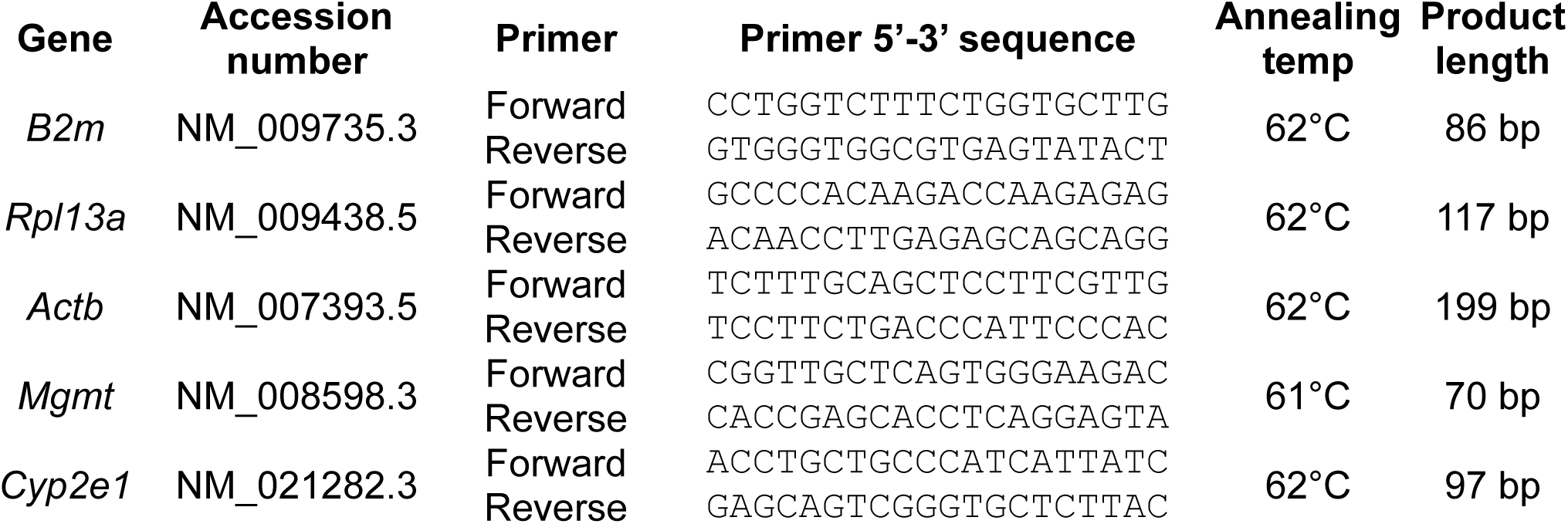
Sequences of primers.

From the three reference genes (*B2m, Rpl13a and Actb*), the least intra-group and inter-group dispersion was observed for *B2m*, so this gene was selected as a reference for all qPCRs (Table S6). Analysis of the resulting qPCR data included averaging technical replicates, dividing the values of target genes to the respective values of the *B2m* reference gene and normalization of all experimental points by the averaged control value. Samples falling outside the calibration range were not counted for analysis, and plates with amplified blanks were rerun. Some outliers determined by boxplots were removed prior to plotting the data and statistical treatment. QPCR data analysis, statistical treatment and visualization were performed using the R package tidyr (50). Data on the plots represent mean ± sd, with the mean of the control group taken as 100%.

### Measurement of MGMT and CYP2E1 protein abundance in mouse livers

Sample preparation and ELISA assays were performed following the instructions from the manufacturers (AVIVA MGMT ELISA Kit Mouse, Aviva Systems Biology, San Diego, CA, cat# OKEH05264; MyBioSource Mouse Cytochrome P450 2E1 (CYP2E1) ELISA Kit, MyBioSource, San Diego, CA, cat# MBS453581). Briefly, ∼100 mg pieces of liver from males and females that had been treated with 5 ppm NDMA for three or ten weeks with respective controls (n=4) were rinsed in 700 μL of 1x cold PBS and homogenized in 1 mL of cold 1x PBS using Dounce homogenizers. Homogenates were stored at −20°C overnight and subsequently subjected to two freeze-thaw cycles in dry ice and in an 8°C water bath, respectively. Cell debris was collected by centrifugation at 5000 x g, 4°C, for 5 min. Based on pilot experiments, supernatants for the MGMT assay were diluted 150 times with Sample Diluent supplied in the kit, while supernatants for CYP2E1 were diluted 225 times with PBS. Both assays were performed immediately after supernatant dilution following instructions from the manufacturers. The samples were assayed in two technical replicates, incubated with peroxidase substrate for 15 min and the results read on the SpectraMax M2e Spectrophotometer (Molecular Devices, San Jose, CA) at two wavelengths: 450 nm (monitoring the peroxidase reaction) and 540 nm (monitoring hemoglobin). Readings at 540 nm were subtracted from the readings at 450 nm to correct for hemoglobin contamination. Raw data and the results, calculated using an R script and linear equation, are presented in Table S7. Data on the plots represent mean ± sd, with means of the control groups taken as 100%.

### Data analysis, statistical treatment and visualization

Data processing, statistical treatment and visualization were performed using the R package tidyr (version 1.3.0) (50). Statistical treatment was performed using an unpaired t-test, with p<0.05 taken as the significance threshold. Bonferroni correction was used to adjust the p value for multiple comparisons. Drawings were made with CorelDraw, Microsoft Excel and BioRender.

## Results

### Setting the dose of NDMA to be used in biochemical and genetic studies

Humans are exposed to NDMA from a variety of sources, by a variety of routes, and at widely different concentrations. While most investigations focus on the biological effects of an acute intraperitoneal administration (3, 31, 47, 51), there are comparatively few studies on chronic low-dose administration of NDMA to experimental animals in drinking water (27, 52) despite the human relevance of this exposure route (10). Recent advances in both highly accurate and sensitive technologies for DNA sequencing and DNA adduct analysis provide opportunities to garner deep insight into the interplay of DNA adducts, DNA repair systems and mutagenic processes in the biological responses to NDMA (31), as well as other carcinogens (53–55). The present study used modern adductomics to measure the biologically relevant DNA adducts of NDMA in liver and used high-resolution sequencing of DNA post-exposure to reveal the distinctive mutational pattern of the carcinogen across all 96 trinucleotide contexts. As described below, these technologies were used in concert to highlight 5 ppm as a dose of NDMA that gave easily detectable adduct and mutation data.

Stable isotope dilution mass spectrometry offers the sensitivity and selectivity needed to monitor DNA adducts in living cells (53, 55–57). This method uses heavy isotope-labeled DNA adducts as internal standards. Heavy DNA adducts are costly to produce by total synthesis so, in the present study, we used a work-around in which *E. coli* cells were grown in the presence of the readily available ^15^N-derivative of ammonium chloride (Figure 3A). The natural biochemical pathways of this microbe utilized the heavy nitrogen to synthesize biomolecules, including DNA. Multi-milligram amounts of ^15^N-enriched DNA were thus produced in bacteria, purified, and then allowed to react *in vitro* with the S_N_1 methylating agent MNU. MNU forms the same DNA adducts as NDMA – specifically, m6G, m7G and m3A (Figure 3B). An LC-MS method was developed that allowed quantitative determination of these three DNA adducts (Figure 3C&D).

Until recently, the lowest known doses of NDMA in chronic regimens in which adducts were quantified in mice were 10 ppm in drinking water (52) or 1 mg/kg with daily intraperitoneal injection (51). The doses used in those studies were dictated by the limits of detection and are high compared to doses typical of human consumption in NDMA-contaminated drinking water and pharmaceutical products. Because current state-of-the-art analytical methods permit improved measurements of chemical and biological effects, our first goal in the present work was to define the dose at which NDMA’s distinctive mutational spectrum emerged above background. This experiment included the administration of NDMA in drinking water at escalating doses and measurement of both hepatic DNA adducts and mutations (Figures 4A, B & D). The abundant adduct, m7G, which represents ∼70% of all damage and is not eliminated quickly by known repair systems (58), served as a marker approximating total alkylation at any given dose. The level of m7G rose nearly linearly from 3 to 40 fmol/µg DNA as the dose of NDMA in water rose from 0 to 2.5 ppm, after which the adduct level plateaued (Figure 4A). The level of the mutagenic adduct m6G in DNA remained at background (∼0.3 fmol/µg DNA) from 0 up to a dose of 1 ppm NDMA; above that dose, the level of adduct rose sharply to 1.5 fmol/µg DNA (Figure 4B).

**Figure 4.**
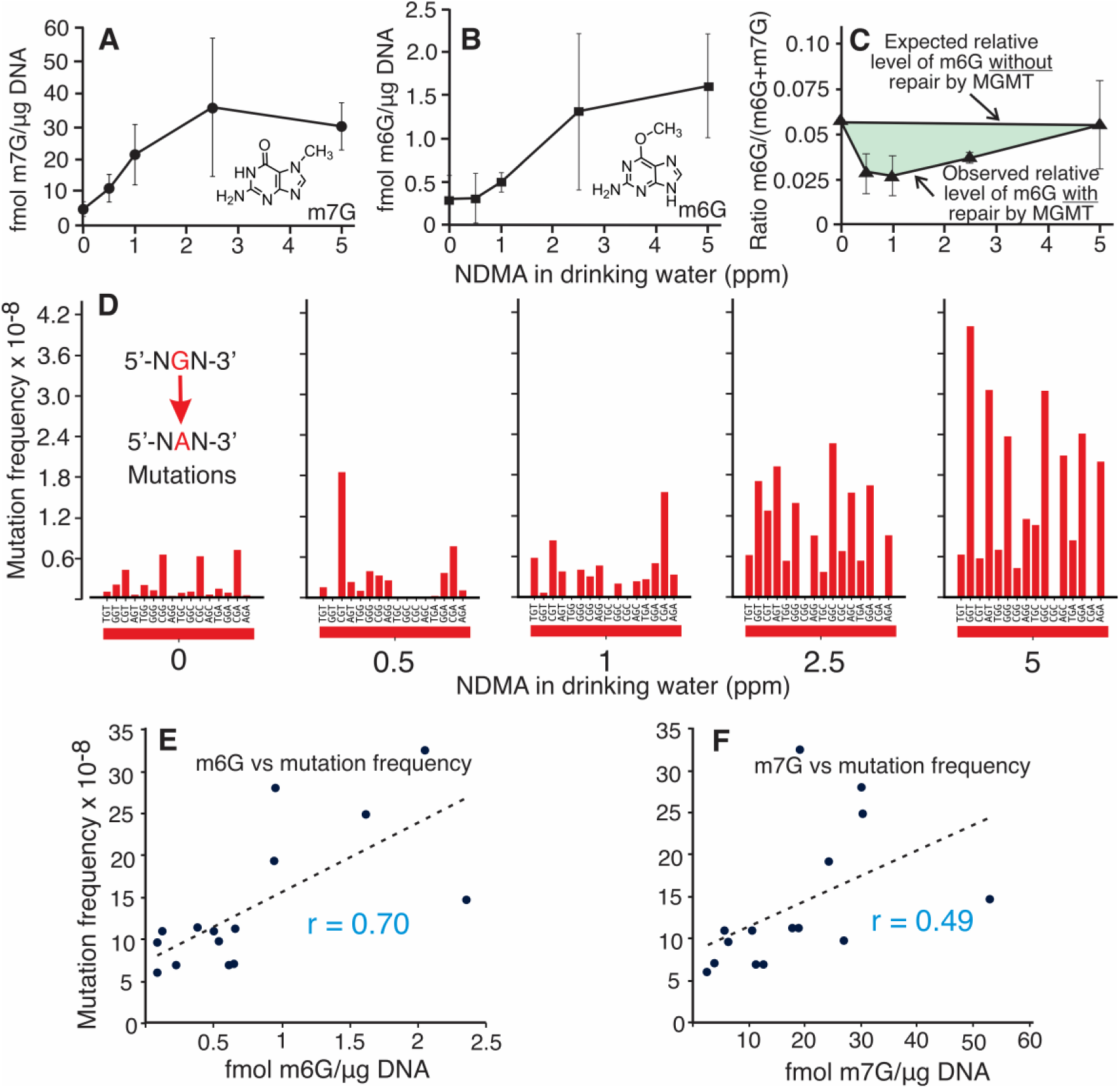
Formation of adducts and mutations in mouse livers. Adducts were measured after two weeks of exposure to NDMA in drinking water at doses ranging from 0-5 ppm, as indicated. **A.** Levels of 7-methylguanine (m7G) in femtomoles of adduct per microgram of DNA analyzed (n=3 for each data point; error bars denote standard deviations). **B.** Levels of O6-methylguanine (m6G) in femtomoles of adduct per microgram of DNA analyzed (n=3 for each data point; error bars denote standard deviations). **C. R**elative levels of m6G as a fraction of the total amount of guanine adducts (sum of the levels of m7G and m6G). In the absence of MGMT repair, it is expected that the relative level of m6G would be independent of dose (the horizontal line). The actual levels observed are below this line, with the shaded area representing the decrease in m6G relative levels that could be attributed to MGMT activity. **D.** The patterns and absolute levels of GC→AT mutations as a function of NDMA dose. The G>A bars were extracted from averaged, normalized and background-subtracted mutational spectra corresponding to each of the NDMA doses. The height of the bars was then normalized to the average G>A mutational frequencies (calculated from sequencing data) for that concentration. **E**. Correlation plot between mutational frequency and levels of m6G adducts. **F**. Correlation plot between mutational frequency and levels of m7G adducts; r denotes the Pearson correlation coefficient.

The inflection point above 1 ppm in the curve depicting m6G vs. dose (Figure 4C) likely signaled the loss of active MGMT, coinciding with an overall increase in the mutational burden, driven primarily by an increase in the GC→AT mutation domain (Figure 4D). It is important to note that the pattern of the GC→AT mutations changes with dose. At 0, 0.5 and 1 ppm, mutations in the GC→AT panels (red bars in Figure 4D) were either random or at 5’-CpG-3’ sites; the latter are typical background mutations from 5-methylcytosine deamination (Figures 4D and Figure S4). It is noteworthy that above 1 ppm, the ability of MGMT to protect genomic integrity waned (Figure 4D), and the sawtooth-like mutation pattern characteristic of m6G emerged. The sawtooth mutational patterns at 2.5 and 5 ppm (Figures 4D and Figure S4) reflect those reported in previous studies after mice or cells are treated with S_N_1 alkylating agents, including NDMA (31, 33, 34, 59).

DNA adducts are well-established precursors to the point mutations characteristic of alkylating agents. It was thus expected that the level of adducts would correlate with the mutational frequency determined by sequencing from the same liver sample. Indeed, correlation plots (Figure 4E and 4F) indicated that the point mutation frequency is positively correlated with the levels of both m7G and m6G. However, the correlation was much stronger (Pearson coefficient r=0.7) in the case of m6G, reinforcing the notion that m6G is the DNA adduct driving the mutagenic outcome of NDMA (Figure 4F).

Another way to visualize the emergence of the mutagenic effect of increasing doses of NDMA is to look at the pattern of point mutations across all DNA sequence contexts. There are six possible types of base substitution mutations, and the breakdown for the NDMA spectra is shown in Figure 5A. While GC→AT mutations comprise a significant proportion even in the control spectra and at low doses of NDMA, they increase considerably at higher doses of the toxicant. This result is consistent with the established notion that a repair threshold exists for m6G, the dominant mutagenic lesion induced by NDMA. We have previously shown that loss of MGMT, the DNA repair protein that processes m6G, results in a large increase in GC→AT mutations (31). Moreover, when comparing the pairwise cosine similarities of the mutational spectra corresponding to the different doses of NDMA (Figure S4), we find that the spectra of controls and NDMA at 0.5 ppm and 1 ppm cluster together (Figure 5B). At the same time, the mutational spectrum at the 5 ppm dose of NDMA clusters closely with COSMIC mutational signature SBS11, a mutational pattern computationally extracted from sequenced human cancer genomes, which we and others have shown to reflect m6G-induced mutagenesis (31, 60). The pattern corresponding to NDMA at 2.5 ppm, while it has some features of the control, clearly shows the emergence of an SBS11-like spectrum. Taken together, the sawtooth-like mutation pattern and corresponding DNA adduct data in Figures 4 and S4 indicate that the 2.5 ppm dose of NDMA, under the conditions of this experiment, is near or above the threshold at which MGMT repair capacity is saturated, and thus the m6G mutational effects are becoming apparent. And, at 5 ppm, a fully mature sawtooth pattern dominates the mutational spectrum.

**Figure 5.**
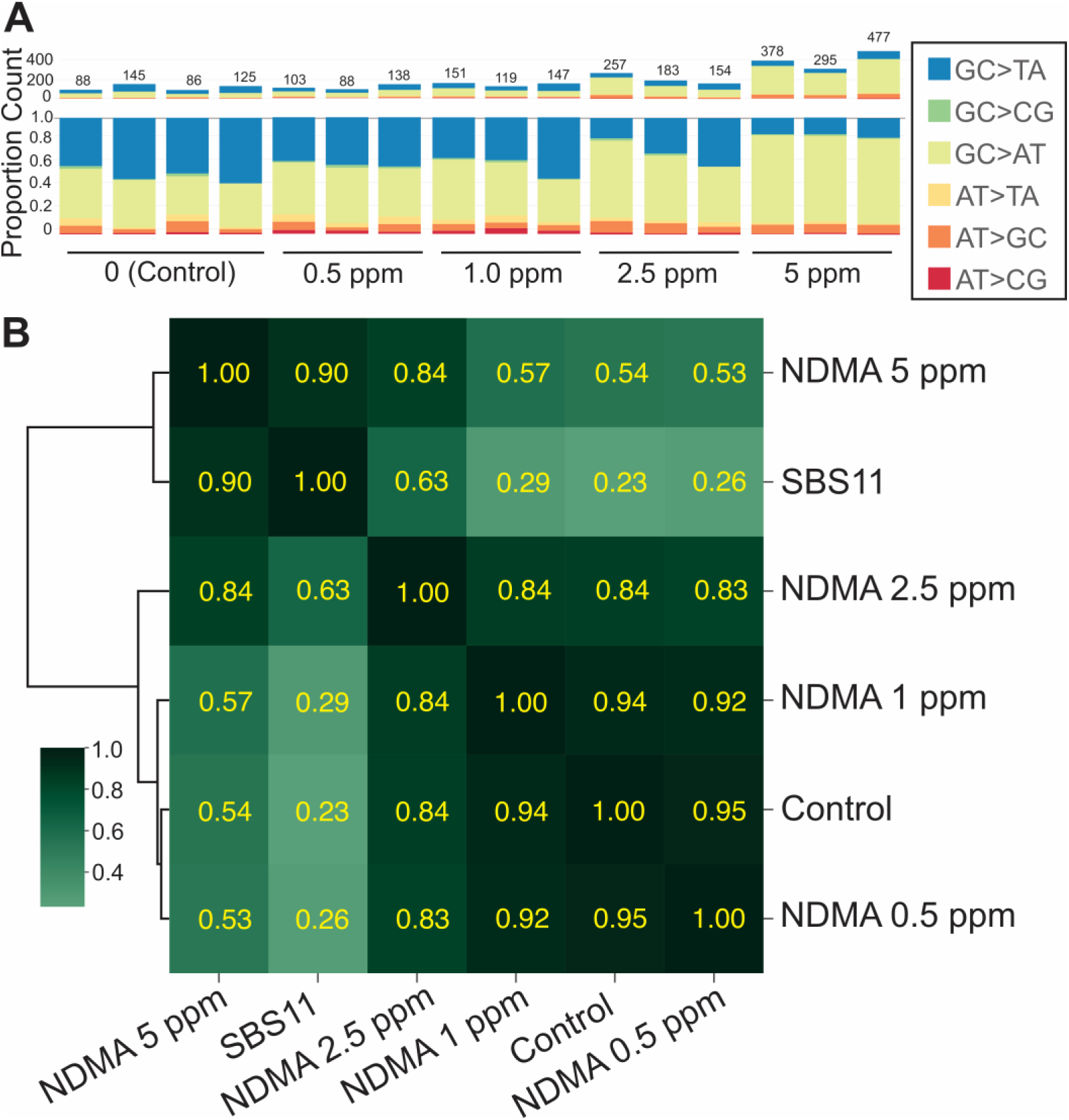
Comparison of mutational spectra corresponding to different doses of NDMA in drinking water. **A.** Colormap of the point mutation distribution across the 6 kinds of point mutations, for individual biological replicates. The numbers above the bars indicate the total number of point mutations identified by duplex sequencing. **B.** Cosine similarity matrix for the mutational spectra corresponding to 2-week exposures to NDMA in drinking water at the levels indicated. COSMIC mutational signature SBS11 is also included for reference.

### Adducts and mutations in male and female mice dosed with 5 ppm NDMA in drinking water

The results of dose-response experiments described here led to the selection of 5 ppm NDMA for the remaining genetic and biochemical studies. After three weeks of exposure to this dose, the levels of m6G and m7G in livers were well above the control level (p < 0.05), whereas the levels of m3A were not significantly elevated over the natural background (Figure 6A). It was of interest to compare adducts and mutations in the livers of male and female mice because males are often more sensitive than females to the toxic and carcinogenic effects of toxicants, including NDMA (61). Contrary to this expectation, females had a higher load of the mutagenic DNA adduct m6G than males after three weeks of dosing. The levels of m3A and m7G were not significantly different between males and females (Figure 6A). As expected, we noted that there was a background level of all three adducts in DNA, likely owing to the presence a ubiquitous endogenous alkylating agent, presumably S-adenosylmethionine or an as-yet uncharacterized nitroso compound (62, 63).

**Figure 6.**
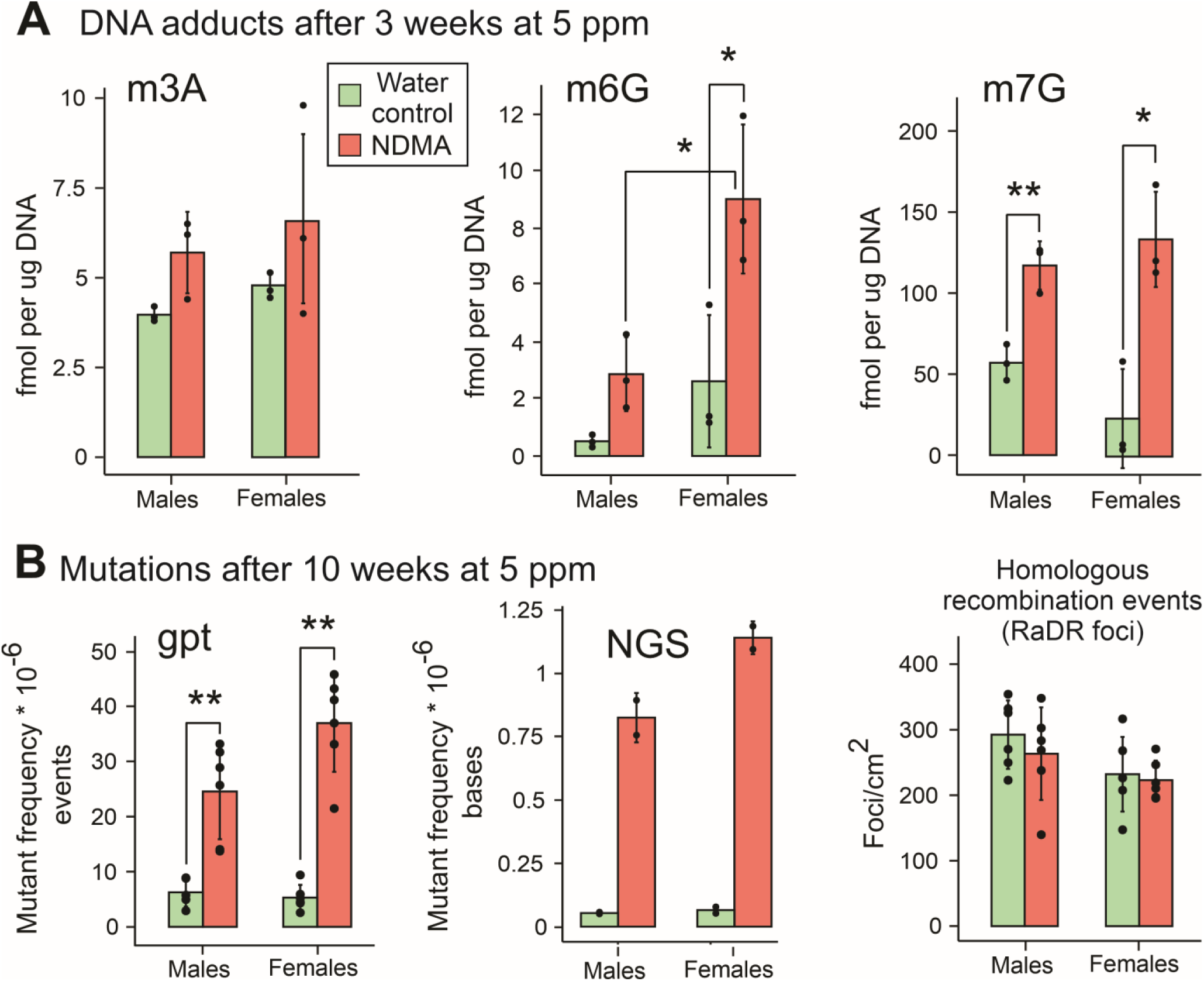
DNA adducts and mutational burden induced by NDMA administered in drinking water. **A.** Alkylation DNA adducts m3A, m6G and m7G were measured in DNA isolated from the livers of mice exposed to 5 ppm NDMA in drinking water for three weeks. **B.** Mutation frequency in the livers of mice exposed to 5 ppm NDMA in drinking water for ten weeks was estimated by the *gpt* assay (left), duplex sequencing (middle) and the *RaDR* assay (right). Height of bars indicates the mean of each distribution, with error bars denoting 1 standard deviation. Statistics: \**p* < 0.05, \*\**p* < 0.01, \*\*\**p* < 0.001 determined by the *t* test. Bonferroni correction was applied for multiple comparisons.

While DNA adducts were measured at three weeks on the 5-ppm regimen, mutations were assayed at ten weeks to allow the adducts sufficient time to be processed into their downstream mutations (Figure 6B). Point mutations (mainly single base-pair substitutions) were measured by two techniques. The *gpt* assay, which measures mutations in the 459 bp *guanine phosphoribosyltransferase* gene, revealed a 3- to 5-fold increase in mutations in livers of NDMA-treated mice (*p* < 0.01); there was no statistical difference between males and females, although there was a trend toward more mutations in females. If accurate, that trend would correlate with the DNA adduct data showing an excess of m6G in females (Figure 6B). The same trend-favoring mutations in females was also observed with a second assay that used next-generation sequencing to measure the mutation rate in the much larger murine genomic domain analyzed by duplex sequencing using TwinStrand hybrid capture probes. This method queries 20 genome loci and a total of 48,000 nucleotides in the mouse genome, with a sequencing depth of approximately 30,000. This method revealed a striking 10- to 15-fold increase in mutant frequency over the control after NDMA treatment. Once again, there were hints of an excess of mutations in females (Figure 6B), but the differences seen were not statistically significant. A third tool used to measure mutations took advantage of the *RaDR* construct genetically engineered into the mouse used in this study. This assay monitors homologous recombination-driven large-scale genomic rearrangements induced by mutagens (47). The *RaDR* assay did not reveal an elevated mutation rate in NDMA-dosed animals, in either sex, at the 5 ppm dose evaluated (Figure 6B, Figure S3).

### The high-resolution mutational spectrum of NDMA reveals a striking sequence context specificity

As indicated above, the collection of mutations from duplex sequencing can be plotted as a function of the trinucleotide contexts, 5’-NXN-3’, in which they occur. (The N residues are normal bases flanking the mutant base, X). The mutant trinucleotide contexts are enumerated to construct a high-resolution mutational spectrum, or HRMS. Although not always done in the literature, it is a good practice in studies such as this one to normalize the raw sequence data by accounting for the frequency of each trinucleotide context in the mouse genome. Some contexts are under- or over-represented, and this normalization puts all contexts on an equal footing as targets for sequence-specific DNA damage, repair and mutagenic polymerase bypass (60). The HRMS of NDMA in mouse liver dosed for 10 weeks at 5 ppm is shown in Figure 7B. The control in which water without NDMA was provided to the mice is shown in Figure 7A. The most notable feature in the NDMA-induced spectrum is the preference for adducts, most likely m6G, to cause mutations in 5’-purine-<u>G</u>-3’ contexts (where <u>G</u> is the position of the mutation). The GC→AT mutations are designated by red bars in the figure. As reported above, this pattern resembles SBS11 seen in several human cancers (64) and is emerging as an exposure biomarker for S_N_1-acting DNA methylating agents (31, 59). The second most dominant mutation type in Figure 7B was the AT→GC transition, which occurred primarily in 5’-TAC-3’ and 5’-GAC-3’ contexts. This pattern resembles a mutational signature called SBS119, which is seen in some human neural cancers treated with DNA methylating agents (65, 66). A third type of mutation, AT→TA, was also observed, comprising only ∼2% of the total mutational burden.

**Figure 7.**
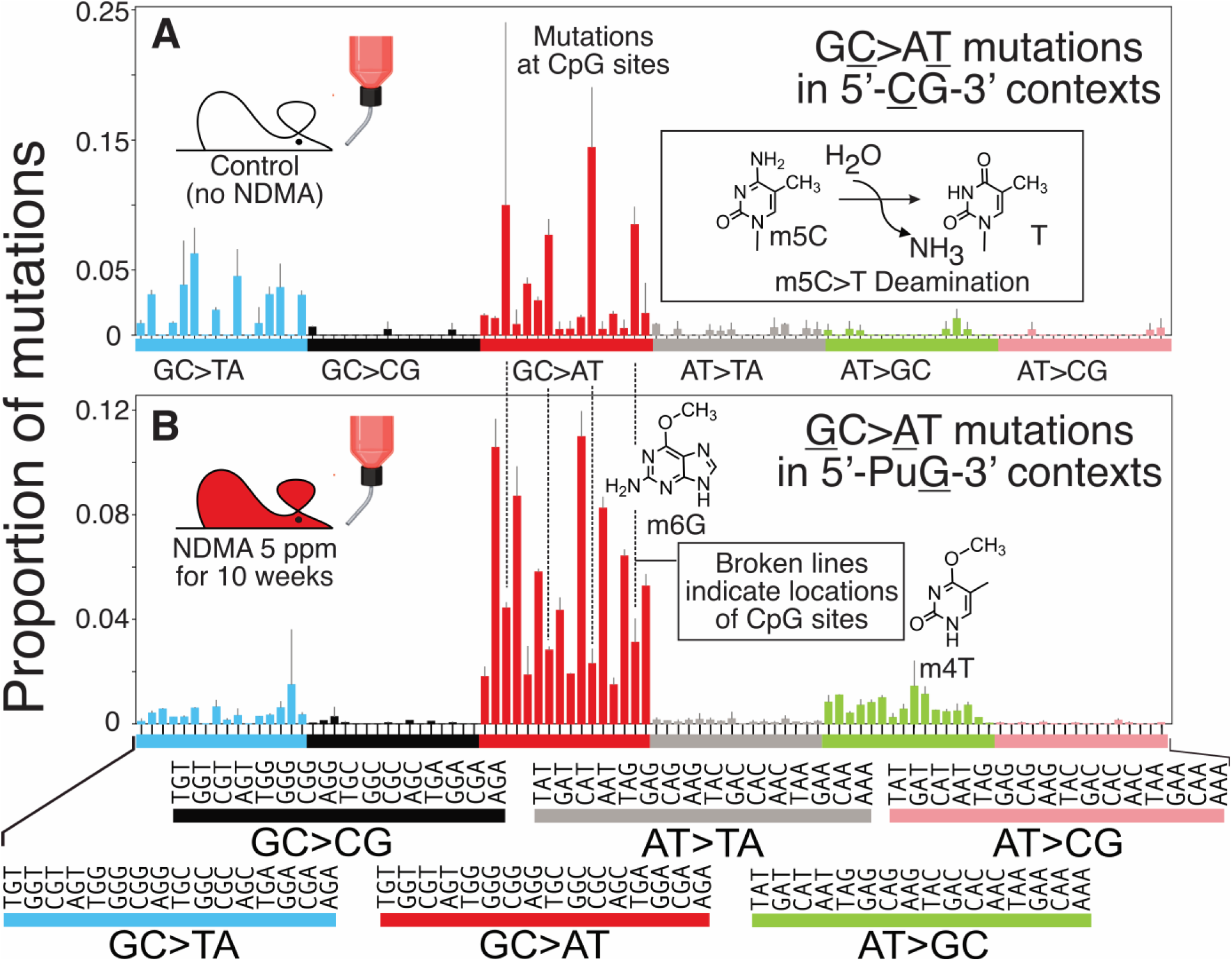
High-resolution mutational spectra in mouse livers after 10 weeks of exposure to 5 ppm NDMA in drinking water. **A.** Mutational spectrum recorded from control animals, receiving no NDMA in drinking water. **B.** Mutational spectrum recorded from animals exposed to 5 ppm NDMA in drinking water for 10 weeks. The spectra are averages of 6 different mice, normalized per-trinucleotide. Error bars denote standard deviations.

### Evaluation of the roles of CYP2E1 and MGMT in cellular responses to NDMA

The elevated level of m6G in female mice seen in Figure 6 prompted an experiment to determine if the higher observed level of adduct was due to preferential metabolism of an electrophile in females. CYP2E1 (the product of the *Cyp2e1* gene) is the principal metabolic enzyme that transforms NDMA into its DNA-damaging electrophilic form (67) (Figure 8A). Examination of this metabolic activation protein was warranted because some previous studies point to the possible role of CYP2E1 in sex-, or sex hormone-mediated conversion of toxicants to biologically active forms (68). Our data, however, did not show sex-specific differences between CYP2E1 mRNA or protein levels at the two terminal timepoints of the experiment: three weeks and 10 weeks (Figure 8 B-E; Figure S5A&B).

**Figure 8.**
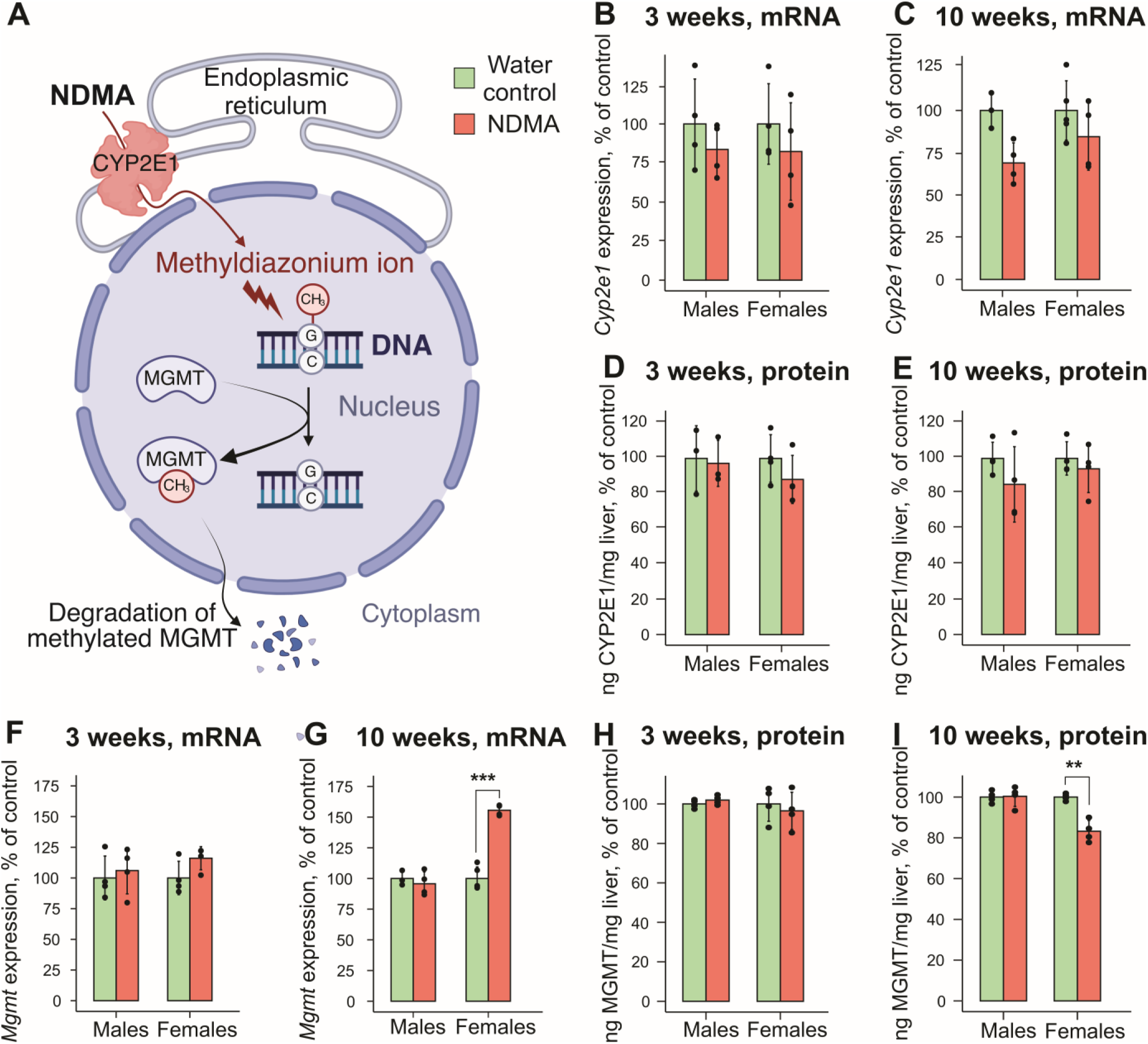
Expression of *Cyp2e1* and *Mgmt* in mouse livers after three and ten weeks of exposure to 5 pm NDMA. **A.** Scheme of CYP2E1 and MGMT contributions to NDMA-mediated mutagenesis. **B, C**. mRNA levels of CYP2E1 in male and female livers after three and ten weeks of exposure to 5 ppm NDMA. **D, E**. protein levels of CYP2E1 in male and female livers after three and ten weeks of exposure to 5 ppm NDMA. **F, G**. mRNA levels of MGMT in male and female livers after three and ten weeks of exposure to 5 ppm NDMA. **H, I.** protein levels of MGMT in male and female livers after three and ten weeks of exposure to 5 ppm NDMA. Statistics: \**p* < 0.05, \*\**p* < 0.01, \*\*\**p* < 0.001 determined by the *t* test.

Another possible explanation of the higher levels of m6G in females could be that the adduct preferentially accumulates and is retained in the genomes of females, possibly owing to reduced DNA repair capacity of the cell’s MGMT reserves. As indicated above, MGMT (the product of the *Mgmt* gene) is the protein that removes the methyl group from m6G, erasing the mutagenic property of the DNA adduct (Figure 8A). Figure 8G shows that females have higher levels of expression of the *Mgmt* gene than males, but they have less of its gene product, the MGMT protein (Figure 8I). These observations were made after 10 weeks on the NDMA regimen; no differences in MGMT expression levels between males and females were observed with the 3-week regimen. These observations can be rationalized by assuming that m6G levels, which are higher in females (Figure 6A), elicit a toxic response in the 10-week regimen. Toxicity implies that the MGMT protein levels are largely depleted (Figure 8I), and thus the need to produce additional MGMT manifests as an increased level of expression of the *Mgmt* gene. As noted, this result is only seen in the 10-week exposure regimen, suggesting that the shorter 3-week regimen did not generate the same level of toxicity to trigger the transcriptional upregulation of the *Mgmt* gene (Figure 8F, Figure S5 C&D).

## Discussion

A key feature of the present work was the coordinated measurement of DNA adducts and mutation frequencies, which revealed mechanistic insights into the ways by which a dosing regimen and sex affect the ability of an animal to respond to DNA damage. Further, the study defined the point at which cellular defenses against the toxicant NDMA became saturated. Significant differences were observed between the mutagenic responses of young mice (>6 weeks old) to NDMA using the chronic dosing model presented here, compared to our previous work (31, 47) in which the toxicant was given to neonates (1-2 weeks old) by acute administration. In our earlier work, a 10.5 mg/kg dosing regimen in which NDMA was administered i.p. to neonatal male and female mice resulted in 45-fold and 30-fold increases, respectively, in *gpt* mutation frequency above background (31). In the present work with chronic NDMA administration, a total dose of 56 mg/kg given over 10 weeks starting at postnatal day 45 produced only 5- and 7-fold enhanced *gpt* mutation frequencies over background in males and females, respectively. Taken together, the data show that a lower dose rate results in disproportionally fewer mutations than an acute dosing regimen, even when the chronic regimen delivers a higher total amount of NDMA. Several factors may explain these observations. First, NDMA pharmacodynamics may differ between the two dosing regimens. Second, NDMA may be more mutagenic in the previously employed neonatal model, where it was given by acute administration, because of the more rapid rate of cellular proliferation in neonates relative to that of young adults. Lastly, and perhaps most importantly, the delivery of NDMA at a low dose rate, as is characteristic of oral administration, may allow for more effective repair of DNA adducts.

The biphasic pattern of m6G accumulation in liver DNA was typical of an adduct for which there is a saturable DNA repair system, in this case, MGMT. MGMT is a suicide protein that is inactivated by the irreversible direct transfer of the methyl group from m6G to its Cys-145 amino acid in its active site, followed by proteasomal degradation of thereby inactivated MGMT (69). For NDMA doses less than 1 ppm (which is the inflection point in the curve in Figure 4B), MGMT is highly effective in preventing NDMA-induced mutagenesis. This is evidenced by an imperceptible increase of m6G adduct levels with dose and a corresponding insignificant change in mutational load (Figure 4B and 4D). Moreover, the mutational patterns corresponding to doses up to 1 ppm show essentially no contribution from SBS11. However, at doses above 1 ppm, both the m6G adduct levels and overall mutational load increase rapidly with dose, while mutational patterns acquire progressively more SBS11-like character. These changes suggest that above 1 ppm, MGMT repair capacity has been exceeded. At the threshold dose of 1 ppm, there was approximately one m6G per 1.5×10^6^ nucleotides, or about 2,000 m6G per genome. Accordingly, in this experimental system, MGMT activity was saturated in mouse livers at NDMA doses that produce ∼2,000 m6G molecules, or more. At these doses, the rate of mutagenic DNA adduct formation exceeds the rate of its removal by MGMT.

One surprising finding in the present study was the effect of sex on the levels of adducts and mutations. Counter to expectations, females had more of the mutagenic DNA adduct m6G than males after three weeks of dosing, even though the levels of m3A and m7G were not significantly different between the sexes. Examination of the expression of the *Cyp2e1* gene and the amount of its encoded protein, CYP2E1, did not adequately explain the higher level of m6G in females, as compared to males (Figures 8B-E). A second possibility was that MGMT levels may differ between sexes during the treatment regimen. In both sexes, there was no change in Mgmt mRNA and protein content after exposure to 5 ppm NDMA for three weeks (Figure 8F&H). After ten weeks, however, Mgmt mRNA was strikingly increased in females but not in males (Figure 8G); this result could reflect a sex-specific compensatory response to the higher level of m6G experienced by females (Figure 6A). Also at ten weeks, there was a lower concentration of the MGMT protein in females (Figure 8I), which again may reflect selective pressure on the MGMT pool, owing to a higher m6G challenge to the genome. The model that emerges based on these findings is that a three-week exposure regimen did not lower the MGMT stores in either sex to the extent that a geno-protective MGMT-based response was evident. At ten weeks, however, females, which have suffered a higher burden of m6G than males, responded by increasing expression of the gene for MGMT. That model would explain the differential transcriptional responses of male and female mice to NDMA (Figures 8F-I).

Mutational spectra are often strikingly irregular, punctuated by hot and cold spots, because the chemistry of DNA adduct formation, the biochemistry of adduct repair and the process of mispairing during replication are often DNA sequence context-dependent (60). Taken together, those factors mold a highly detailed mutational fingerprint: the high-resolution mutational spectrum (e.g., Figure 7). The rich dataset in the HRMS can be used to search databases for similarities with human diseases and be applied as a diagnostic tool to identify prior exposure to carcinogenic agents (31, 49, 70, 71). The fine-grained details of the HRMS can also be used to (a) nominate specific DNA lesions as the molecular triggers for a given mutation type (60), (b) help define the possible role DNA sequence context and DNA repair systems play in molding the mutational pattern (42), and (c) provide biomarkers that can be used to assess past exposure to a DNA damaging agent (71, 72).

Comparison of the HRMS from acute administration of NDMA published earlier (31) and the present spectra from chronic administration (Figure 7B) show that most mutations are caused by m6G in contexts where a target guanine is preceded on the 5’-side by a purine. These mutations are modulated by MGMT (31) and are presumably caused by the mispairing of m6G with thymine. There are significant differences, however, between the acute and chronic spectra in other aspects of the mutational pattern. To give one example, the fraction of AT→GC mutations is 11% in the genomes of animals exposed to NDMA by chronic oral administration. By contrast, the level of this mutation type is only 5% in mice exposed by acute administration (31). The AT→GC mutations could be caused by the error-prone bypass of adenine- or thymine-derived DNA lesions, including m3A, 1-methyladenine, 3-methylthymine, *O*^2^-methylthymine, and *O*^4^-methylthymine (m4T). While we cannot rule out contributions from the first four adducts, the mutational specificity of m4T (Figure 1B) matches the mutations seen in the AT→GC domain in Figure 7B (73–76). Therefore, our working hypothesis is that the AT→GC mutations are caused by mis-replication of m4T. While much less abundant than m6G, m4T is largely refractory to repair by MGMT and exceptionally mutagenic. Inefficient repair of m4T during chronic exposures would result in m4T accumulation, explaining its potentially high contribution to the mutational burden arising from NDMA exposure. It is noteworthy that some studies of mutagenesis by S_N_1 alkylating agents (e.g., MNU and the chemotherapeutic temozolomide) show a much stronger representation of AT→GC mutations than seen here (71). Those studies were done in cell culture *in vitro*, unlike the studies here, which were done in animals, *in vivo*. One conjecture could be that the *in vitro* studies selected for surviving cells that express high levels of MGMT. That environment would favor repair of m6G, while sparing m4T, leading to a high proportion of AT→GC mutations in the surviving cells in cell culture.

The mutational spectrum of chronically administered NDMA includes, in addition to the aforementioned transition mutations, a small fraction of GC→TA and AT→TA transversions (Figures 5A & 7B). Conventional wisdom suggests that the GC→TA mutations would be due to the formation of 7,8-dihydro-8-oxoguanine (commonly known as 8-oxoguanine or 8-oxoG), which shows this mutational specificity (77). A number of other possibilities exist, however, among the large collection of oxidized guanines (78). The AT→TA mutations in the spectrum could be due to m3A, which is a relatively abundant DNA adduct (Figure 3D); however, the mutagenic consequences of this very toxic DNA adduct have not yet been definitively established. It is also possible that an even rarer adduct, *O*^2^-methylthymine, could contribute to the AT→TA mutations. Site-specific mutagenesis studies show that this poorly repaired adduct does indeed induce T→A transversions (79, 80) when replicated in cells.

With regard to the homologous recombination-driven large scale rearrangement mutations assessed via the *RaDR* assay, an interesting contrast arises between our previous work with an acute dosing regimen to neonates (47) and the current study. The acute regimen previously used shows both abundant NDMA-induced point mutations (via the *gpt* assay) and large-scale gene rearrangements measured by the *RaDR* assay (47). As shown here, however, chronic exposure to NDMA in young adult wild type mice induced point mutations (Figure 4B), but not large-scale genomic rearrangement mutations. Both m3A and m6G can induce strand breaks that stimulate genomic rearrangements (47), but neither lesion seems to be triggering such genetic changes in the present chronic dosing study. To explain this result, we note that the levels of these lesions are relatively low. The combination of low levels of recombinogenic lesions, owing it is assumed to facile repair, and the lower replication rate of young adult mice (as compared to neonates) likely resulted in the absence of detectable genomic rearrangements (as measured by the *RaDR* assay) after chronic low-dose exposure to NDMA.

### Conclusions and perspective

The coordinated measurement of adducts and mutations has practical value in several areas. As one example, m6G is widely regarded as a toxic and carcinogenic threat and understanding the balance between the natural forces that remove it from DNA and the processes by which it is converted into mutations is of paramount importance. As a second point, NDMA is a member of a broad class of DNA alkylating agents that modify DNA by an S_N_1 mechanism. Its DNA adduct profile and corresponding mutational spectrum are similar to that of the chemotherapeutic temozolomide, which is used in regimens that treat glioblastoma multiforme. The tools used herein could be of value to design therapies that produce a sufficient amount of m6G, which is the therapeutic DNA adduct, to trigger a toxic response in the tumor. Tumors often achieve drug resistance by increasing the level of MGMT, so the adductomic and genomic methods presented could allow fine-tuning of a dosing regimen to provide a favorable therapeutic index. Thirdly, while cancer treatment is always important, a lesser studied but also crucial line of research involves finding compounds that can prevent malignancy. Once again, the tools described here could be used to identify agents that intercept the pathways of DNA adduct formation, enhance DNA repair or otherwise reduce carcinogenic risk. Lastly, and perhaps most importantly, the HRMS of NDMA, and of other environmental mutagens (e.g. the hepatocarcinogen aflatoxin B_1_ (72)) typically appear in tissues long before malignant processes become evident. Mutational patterns in tissue or liquid biopsy scenarios could inform early detection of an incipient malignancy and motivate cancer therapy decisions before carcinogenic processes accelerate.

## Acknowledgements

The authors would like to acknowledge and thank the MIT BioMicro Center for DNA sequencing; the MIT Department of Chemistry Instrumentation Core for NMR instrumentation; MIT Center for Environmental Health Sciences bioanalytical core facility for mass spectrometry support; Suzette Morales (MIT 68S animal facility), Maxim Pyatkov for help with data analysis (Boston University), Caroline Atkinson (MIT DCM), Magalie Boucher (MIT DCM), Ed Clark (MIT DCM), Charlie Demurjan for data aggregation and management for FairdomHUB.

## Dedication

The authors dedicate this paper to Thomas W. Kensler (1948–2025), a colleague who pioneered the field of mechanism-informed cancer prevention.

## Data availability

All the data described in this article are available in the main text and figures, in the online supplementary material, and indexed in the FAIRDOM Hub database (https://fairdomhub.org/studies/1397). The sequencing data files have been deposited in the NCBI Sequence Read Archive (SRA) under the BioProject PRJNA1337648 (http://www.ncbi.nlm.nih.gov/bioproject/1337648).

