## Supplemental Figures for "DNA adduct and mutational profiles reveal a threshold of cellular defenses against *N*-nitrosodimethylamine administered to mice in drinking water"

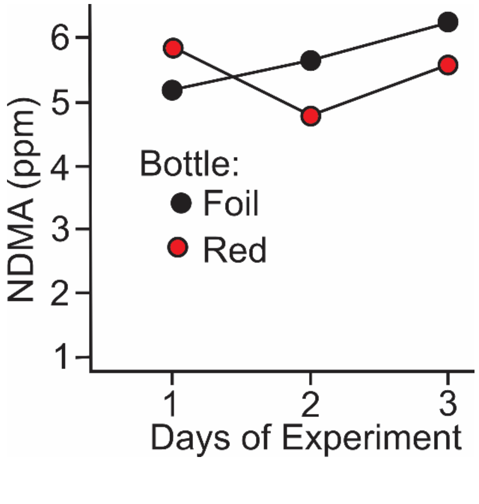


**Figure S1. Stability of NDMA in drinking water.** Solutions of 5 ppm NDMA in water were prepared and aliquoted in both standard red rodent drinking bottles and in clear bottles covered by foil (positive control). The concentration of NDMA was measured daily in both types of bottles. No significant change in the NDMA concentration was detected over the course of 3 days.

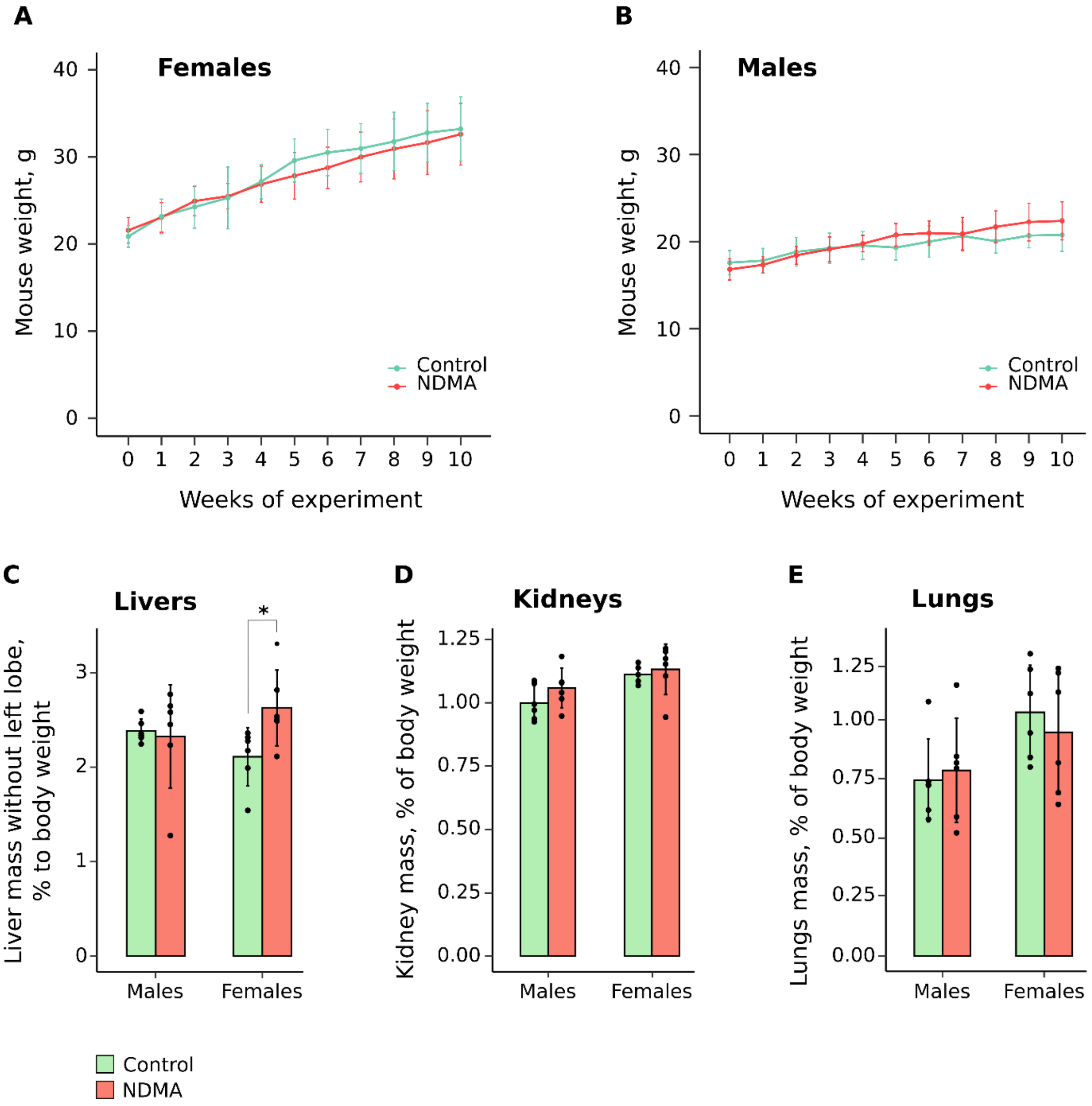


**Figure S2. Mouse body and organ weights after 10 weeks of exposure to 5 ppm NDMA. A**, **B**. Females and males body weights throughout experiment. **C**, **D**, **E**. Weights of livers (without left lobes), kidneys and lungs after 10 weeks of exposure to 5 ppm NDMA. Statistics: *p < 0.05 by the *t* test.

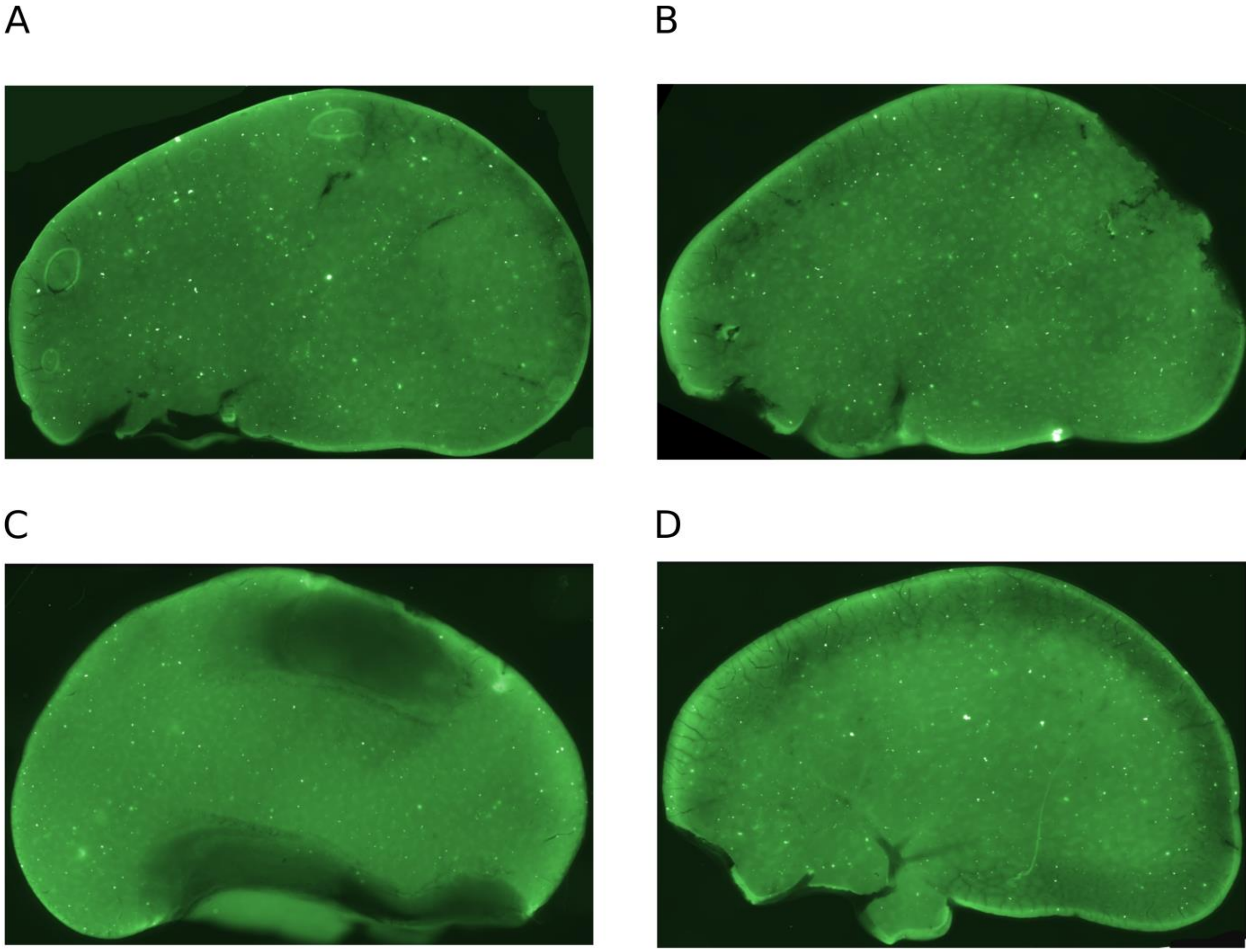


**Figure S3. Representative RADR fluorescent images of left liver lobes after 10 weeks of exposure to 5 ppm NDMA.** Bright dots represent foci of homologous recombination-driven large-scale chromosomal rearrangements. **A**. Male, water control. **B**. Male, 5 ppm NDMA in drinking water. **C**. Female, water control. **D**. Female, 5 ppm NDMA in drinking water. For each condition, n=6 animals were analyzed.

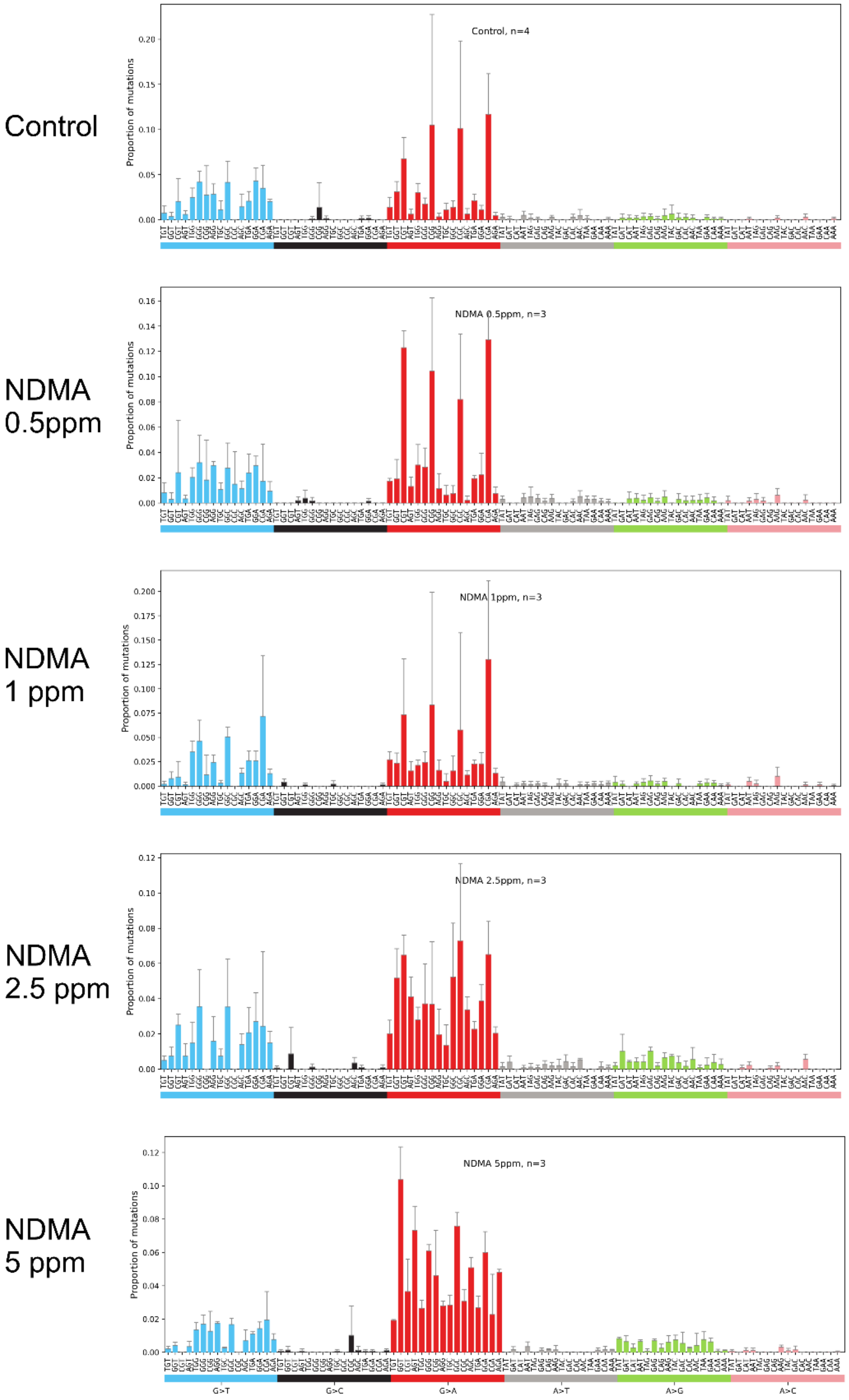


**Figure S4. Mutational spectra in DNA isolated from livers of mice treated with NDMA in drinking water.** Each spectrum is an average of n=3 or n=4 mice, and normalized per-trinucleotide. Height of the bars are means, error bars indicated standard deviations.

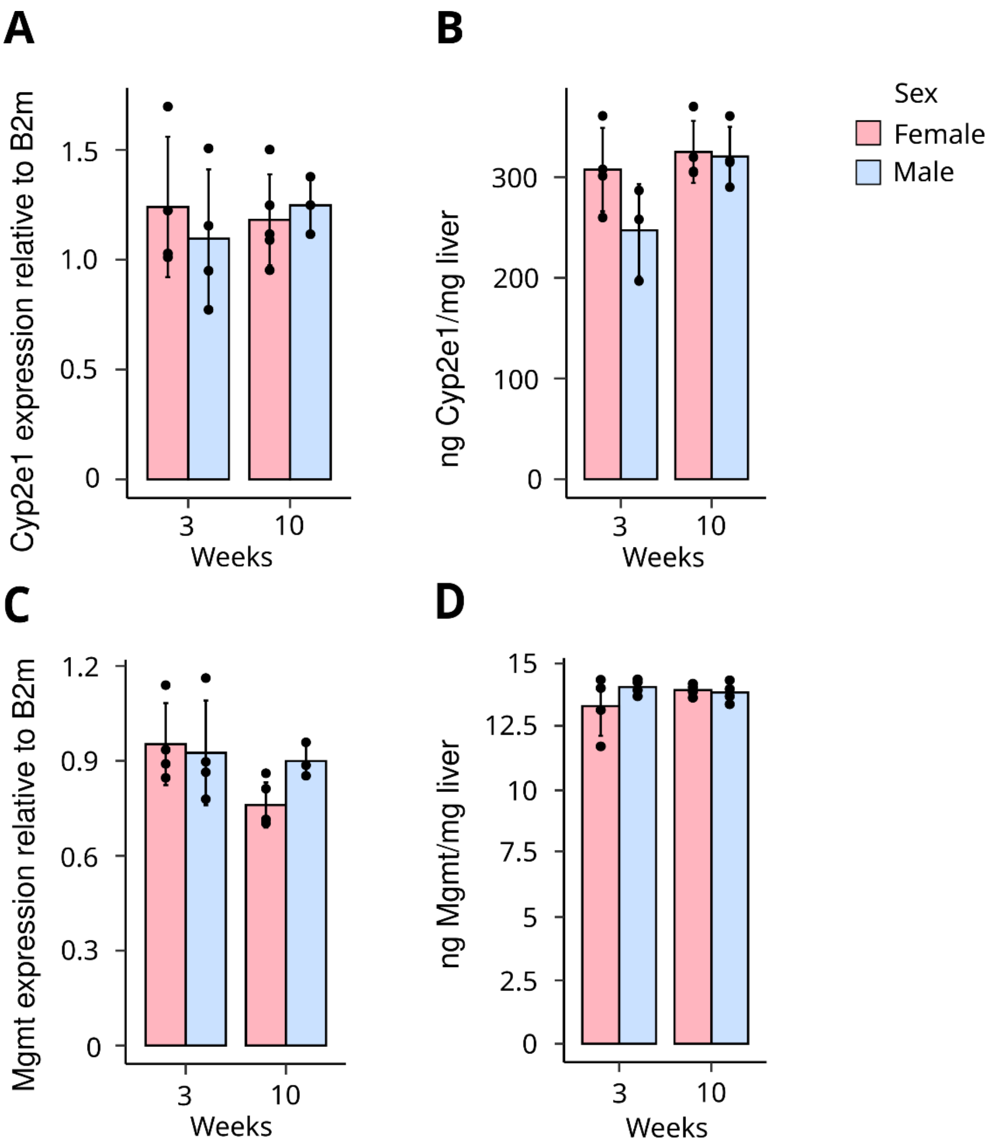


**Figure S5. Relative levels of *Mgmt* and *Cyp2e1* mRNA and protein content in mouse livers after 3 and 10 weeks of exposure to NDMA in drinking water.** **A**. *Mgmt* mRNA levels relative to those of *B2m*. **B**. MGMT protein levels per 1 mg of liver tissue. **C**. *Cyp2e1* mRNA levels relative to those of *B2m*. **D**. CYP2E1 protein levels per 1 mg of liver tissue.
